# The glucocorticoid dexamethasone influences motility of the sulfate-reducing bacterium *Desulfovibrio desulfuricans* by targeting the filament cap protein FliD

**DOI:** 10.64898/2026.07.05.734484

**Authors:** Elena Fajardo-Ruiz, Emanuel Kring, Dominik Schum, Sophie Brameyer, Robin Kretschmer, Tao Wang, Joshua Hesse, Isabella Gantner, Emma Weißert, Lisa Goebner, Lucas Milles, Tobias Gulder, Stephan A. Sieber, Kirsten Jung

## Abstract

Glucocorticoids such as dexamethasone (**DXE**) are first-line treatments for inflammatory bowel disease (IBD). Beyond their effects on the host immune system, accumulating evidence suggests that glucocorticoids can also influence the gut microbiota. Notably, IBD patients exhibit an increased intestinal colonization by sulfate-reducing *Desulfovibrio* spp. Here, we show that **DXE** modulates bacterial motility in the gut commensal *Desulfovibrio desulfuricans* through a metabolism-independent mechanism. To identify bacterial targets, we developed a **DXE**-derived chemical probe and performed affinity-based protein profiling, which revealed the flagellar cap protein FliD (Ddes_0530) as a principal binding partner. Structural modeling using AlphaFold3 and Boltz2 predicted **DXE** binding within a conserved groove of the FliD C-terminal domain. Furthermore, the tip of the flagellum of *Desulfovibrio*, but not that of *Escherichia coli,* could be fluorescently labeled with TAMRA-**DXE**, but not with the structurally related steroid probe TAMRA-norethiosterone, indicating that flagellar labeling is specific to **DXE** rather than the steroid scaffold itself. As a consequence of this interaction, transmission electron microscopy showed that **DXE** treatment prevented flagellation in a subpopulation and reduced flagellar length in *D. desulfuricans* strains ATCC 27774 and CCUG 72978, respectively. Quantitative motility tracking revealed a non-monotonic, dose-dependent modulation of swimming velocity, with peak stimulation at 10 µM **DXE**, accompanied by straighter trajectories and enhanced net displacement. Together, these findings uncover a previously unrecognized mode of action for **DXE** which directly perturbs flagellar biogenesis and motility of an important gut microbiome member of IBD patients.

**Significance:** Glucocorticoids are widely prescribed for inflammatory conditions, yet their direct effects on gut bacteria remain largely unexplored. We demonstrate that dexamethasone, a synthetic glucocorticoid, binds to the flagellar cap protein FliD of the gut commensal *Desulfovibrio desulfuricans*, affecting flagellar assembly and altering motility behavior. Unlike previously characterized steroid-metabolizing bacteria such as *Clostridium steroidoreducens*, *Desulfovibrio* does not degrade dexamethasone, indicating that the observed effects result from direct drug-protein interaction. These findings establish a new paradigm for metabolism-independent drug-microbiome interactions and suggest that glucocorticoid effects on gut bacteria extend beyond enzymatic degradation pathways.

## Introduction

Glucocorticoids are among the most widely prescribed anti-inflammatory drugs, with dexamethasone (**DXE**) serving as a cornerstone therapy for inflammatory bowel disease (IBD), autoimmune disorders and severe infections (1). Despite their clinical importance, patient responses to glucocorticoid therapy vary considerably and the mechanisms underlying this variability remain incompletely understood. Emerging evidence suggests that the gut microbiota may modulate drug efficacy through both metabolic transformation and direct drug-microbe interactions (2–5).

The gut microbiota harbors diverse enzymatic activities capable of transforming xenobiotics, including therapeutic drugs (2). Recently, the OsrABC reductase system was identified in *Clostridium steroidoreducens*, which reduces synthetic glucocorticoids including prednisolone to inactive 3β,5β-tetrahydro metabolites (6). Homologs of this system are enriched in Crohn’s disease patient metagenomes, where they may contribute to glucocorticoid treatment failure (7). However, the metabolic capacity for steroid transformation varies substantially across bacterial taxa and the direct effects of glucocorticoids on bacteria that lack such enzymatic machinery remain largely unexplored (5).

One bacterial genus that lacks known steroid catabolic gene clusters is *Desulfovibrio* (8, 9). These Gram-negative bacteria are characterized by the ability to reduce sulfate to hydrogen sulfide during anaerobic respiration of organic compounds (10) and are widespread in natural environments (11). In humans, *Desulfovibrio* species are naturally present in the gut microbiomes of approximately 50% of the population, and make up less than 1% of the total bacterial population in a healthy host (10–12). A massive proliferation of *Desulfovibrio*, known as a bloom, has been associated with various gastrointestinal conditions and their hydrogen sulfide production may influence gut barrier function and inflammation (13–19).

Besides their metabolic properties, motility is another important determinant of *Desulfovibrio* physiology and host colonization. Flagella-mediated motility plays a crucial role in bacterial host colonization, biofilm formation and pathogenesis (20). The bacterial flagellum is a complex nanomachine composed of a basal body, hook and filament (21). Flagellar assembly requires the coordinated action of numerous proteins, including the filament cap protein FliD, which facilitates the polymerization of flagellin subunits at the growing filament tip (21, 22). FliD forms oligomeric complexes: hexamers in *Escherichia coli*, pentamers in *Salmonella* and tetramers in *Bdellovibrio bacteriovorus* (23–25). These oligomeric formations are essential for efficient flagellar assembly (22, 23). Given the central role of motility in bacterial physiology, increasing attention has been directed toward host-derived molecules that modulate this process.

Interkingdom signaling between eukaryotic hosts and bacteria has been demonstrated for catecholamine hormones, which bind to the chemotaxis coupling protein CheW in *Vibrio campbellii*, affecting bacterial motility and chemotactic control (26). Whether similar direct interactions occur between synthetic drugs and bacterial proteins remains an open question with significant implications for understanding drug-microbiome interactions (3, 4, 27).

To address this question, we investigated whether **DXE** directly interacts with proteins of *D. desulfuricans* and how such interactions influence bacterial physiology. We found that **DXE** alters the swimming behavior of *D. desulfuricans*. We applied chemical proteomics to identify protein targets of **DXE** in *D. desulfuricans*. For this purpose, we designed and synthesized a tailored **DXE** probe and identified the flagellar cap protein FliD as a major target. Structural modeling and in situ labeling supported **DXE** binding to FliD. Functional studies revealed that **DXE** treatment affects flagellar assembly and alters motility behavior. This study reveals a previously unknown effect of glucocorticoids on *Desulfovibrio*, underlining the importance of considering microbiome-drug interactions in IBD therapy.

## Results

### Dexamethasone Alters Swimming Behavior in *D. desulfuricans*

Based on our experiences of the effect of catecholamines on motility (26), we first examined the effect of **DXE** on *D. desulfuricans* bacterial swimming behavior using swim agar assays. These assays were performed in the presence of 10 µM **DXE** using the clinical isolate *D. desulfuricans* CCUG 72978. *D. piger* (a non-motile *Desulfovibrio* species) as well as *E. coli* were used as controls. While we consistently observed a readily detectable phenotype consisting of distinct ring formation patterns in the presence of **DXE** across multiple biological replicates of *D. desulfuricans* CCUG 72978 (Fig. 1*A*), quantitative measurements of the swim diameter did not reveal significant differences (*p* = 0.36, repeated-measures one-way ANOVA, Fig. 1*B*), suggesting that **DXE** alters the pattern of bacterial motility rather than its overall extent. The **DXE**-treated plates exhibited clear, defined concentric rings with a pattern-forming expansion front compared to both control conditions, which showed smooth, uniform spreading without ring segmentation. This qualitative phenotype suggests that **DXE** may influence the temporal dynamics of bacterial swimming or cell density distribution without affecting the overall swim distance. *E. coli* showed no phenotypic changes in response to **DXE** treatment, indicating a specific effect.

**Figure 1.**
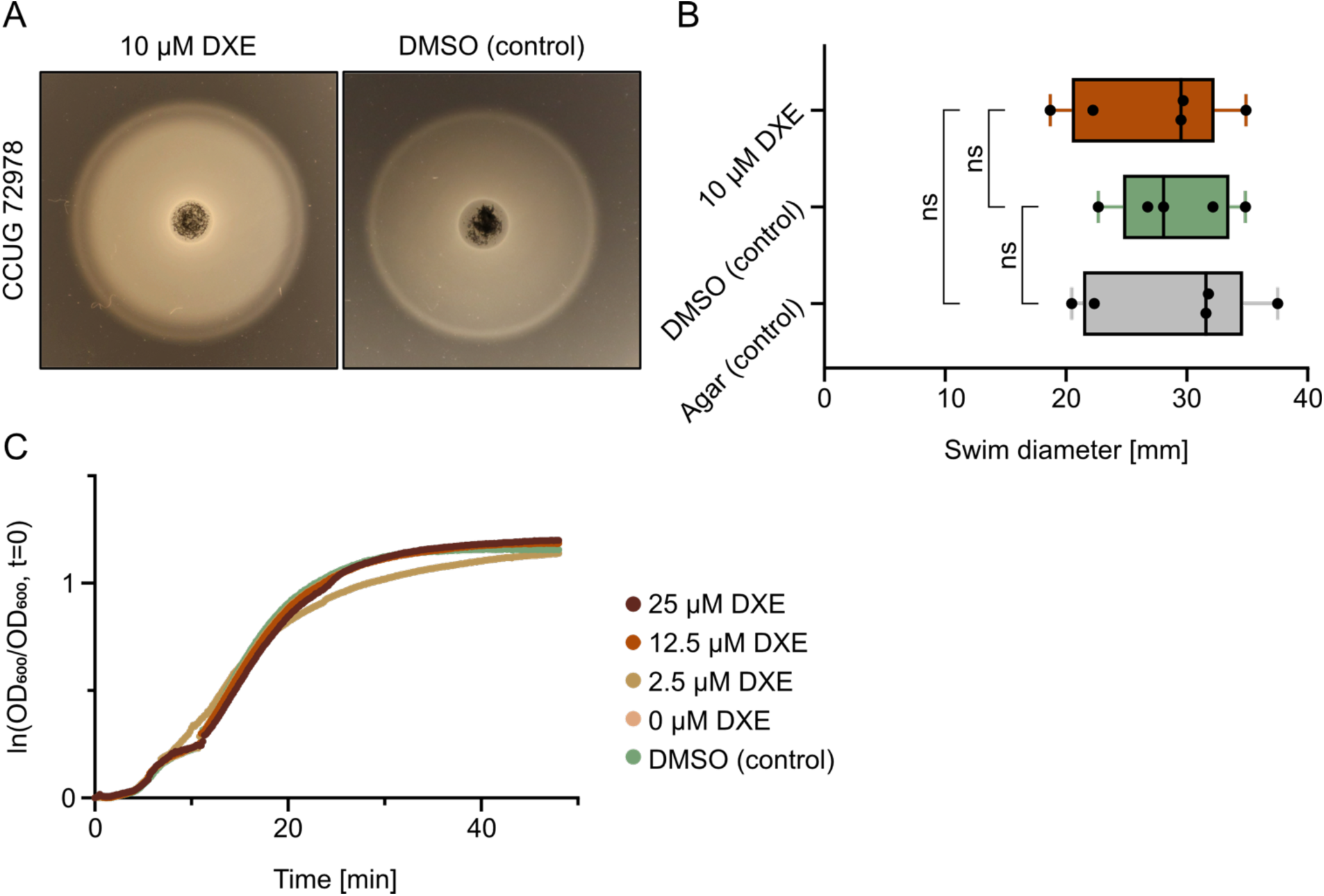
Dexamethasone alters swimming behavior without affecting the growth of *Desulfovibrio*. (*A*) Representative swim agar plates of *D. desulfuricans* CCUG 72978. Bacteria were spotted on 0.3% Columbia soft agar, containing 10 µM dexamethasone (**DXE**) or an equal volume of DMSO (control), and incubated anaerobically at 37 °C for 48 h. (*B*) Quantitative measurements of swim diameter at the indicated conditions (p = 0.36, repeated measures one-way ANOVA; n = 5 biological replicates). Error bars represent SEM; ns, not significant. (*C*) Growth of *D. desulfuricans* CCUG 72978 in Postgate medium, supplemented with 0, 2.5, 12.5, or 25 µM **DXE**, or DMSO vehicle. Growth was monitored by OD_600_ measurements every 10 min for 48 h. Data are presented as mean ± SEM of normalized ln(OD_600_) values (one-way ANOVA: *p* = 0.43 for CCUG 72978; *n* = 3 biological replicates).

### Dexamethasone Does Not Significantly Affect *Desulfovibrio* Growth and is Not Metabolized

To determine whether the observed motility phenotype resulted from growth effects, we performed comprehensive growth studies under anaerobic conditions using three *D. desulfuricans* strains (ATCC 27774, CCUG 72978 and DSM 642). Cultures were treated with **DXE** at 2.5, 12.5 and 25 µM, alongside DMSO vehicle and untreated controls and growth was monitored over 48 hours by OD_600_ measurements in biological triplicates.

All three *D. desulfuricans* strains exhibited robust and consistent growth across all conditions as indicated by the growth of *D. desulfuricans* CCUG 72978 (Fig. 1*C*). **DXE** treatment at concentrations up to 25 µM, encompassing and exceeding typical therapeutic plasma concentrations (0.1 - 1 µM) (28), did not significantly affect maximum culture density compared to DMSO vehicle controls.

Furthermore, we assessed **DXE** stability during anaerobic growth of *D. desulfuricans* ATCC 27774 and CCUG 72978 using liquid chromatography coupled to high-resolution mass spectrometry (LC-HRMS). **DXE** was detected in the protonated form ([M+H]^+^, *m/z* 393.2066) with a retention time of approximately 12 minutes, matching the authentic standard (Fig. 2*A*). Compound identity was confirmed by MS^2^ fragmentation, which yielded diagnostic product ions at *m/z* 147.0795, 237.1264, 279.1733 and 355.1896, consistent with the expected fragmentation pattern of the fluorinated steroid scaffold (Fig. 2*B*). Time-course analysis revealed that **DXE** signal intensity remained stable throughout the five-day incubation period in both *D. desulfuricans* ATCC 27774 and CCUG 72978 (Fig. *2A, C*). Quantitative comparison of extracted ion chromatogram peak areas showed no statistically significant decrease relative to day 0 controls. Furthermore, comprehensive analysis of the full chromatographic profiles revealed no emergence of novel peaks that would indicate the formation of **DXE** metabolites.

**Figure 2.**
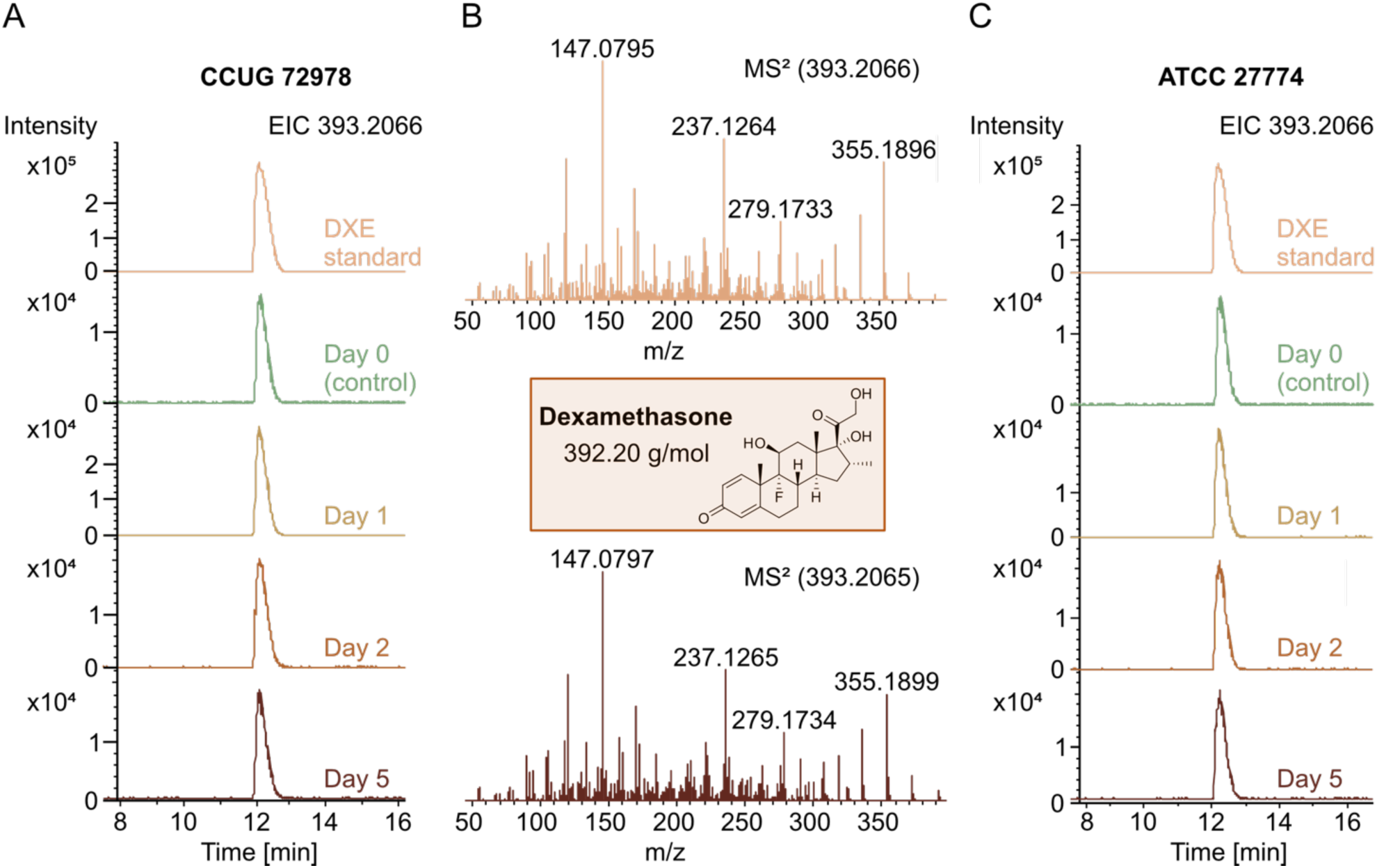
Dexamethasone is metabolically stable in *Desulfovibrio* cultures. (*A*) Extracted ion chromatograms (EIC) of **DXE** ([M+H]^+^, *m/z* 393.2066; mass tolerance ±5 ppm) from culture supernatants of *D. desulfuricans* CCUG 72978 collected over a five-day time course at days 0, 1, 2, 3, and 5. (*B*) High-resolution MS^2^ fragmentation spectrum of the **DXE** peak, acquired by data-dependent acquisition on a Q Exactive HF mass spectrometer (Thermo Fisher Scientific). Diagnostic product ions at *m/z* 147.0795, 237.1264, 279.1733, and 355.1896 confirm the identity of the intact fluorinated steroid scaffold. (*C*) EIC chromatograms from culture supernatants of *D. desulfuricans* ATCC 27774 collected at the same timepoints after anaerobic incubation with 50 µM **DXE** at 37 °C. Quantitative comparison of peak areas revealed no statistically significant decrease relative to day 0 controls for either strain. The **DXE** peak elutes at a retention time of approximately 12 min.

In line with a lack of functional steroid-transforming enzymes such as the OsrABC reductase system found in *C. steroidoreducens* (6), these results demonstrate that *D. desulfuricans* does not metabolize **DXE** under the conditions tested. This finding indicates that any effects of **DXE** on *Desulfovibrio* physiology must result from direct drug-protein interactions, not metabolic transformation.

### Design and Synthesis of a Dexamethasone Probe for Target Identification

To identify the molecular targets of **DXE** in *D. desulfuricans*, we designed and synthesized a chemical probe for protein enrichment via affinity-based protein profiling (A*f*BPP) (Fig. 3*A*). Even though **DXE** contains an α,β-unsaturated carbonyl moiety within the A-ring, this motif is embedded in a conjugated and sterically constrained steroid scaffold. Thus, it is unlikely to function as an efficient intrinsic warhead for covalent protein binding. In addition, **DXE** engages its primary molecular target, the glucocorticoid receptor, through non-covalent binding interactions (29, 30). Building on previous studies that confirmed C21 position of **DXE** to tolerate structurally diverse substituents with comparatively little impact on receptor binding, we selected the 21-hydroxyl group as the derivatization site for a probe (31–33). Based on this, we designed and synthesized a **DXE**-derived photoaffinity probe (*photo*-**DXE**) that retains the core steroid scaffold while introducing two functional elements required for target engagement: a diazirine moiety which serves as a photoactivatable crosslinking group enabling covalent capture of **DXE**-protein interactions upon UV irradiation, and a terminal alkyne handle for downstream copper-catalyzed azide-alkyne cycloaddition (CuAAC; “click chemistry”) allowing conjugation to reporter or affinity tags such as fluorophores for visualization or biotin for protein enrichment.

**Figure 3.**
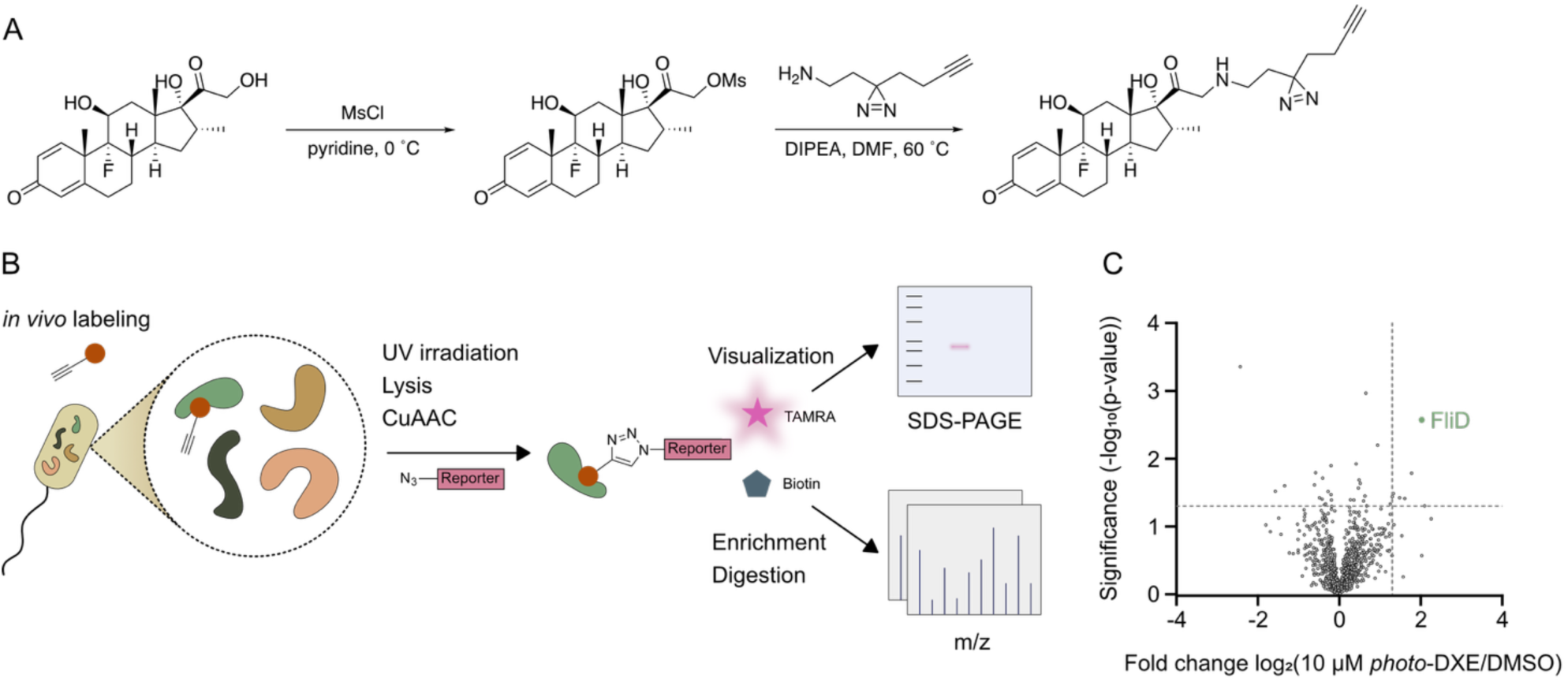
Chemical proteomics identifies FliD as a dexamethasone target. *(A)* Synthetic route for the preparation of the **DXE**-probe starting from dexamethasone (**DXE**). The terminal hydroxyl group of the side chain was first converted into the corresponding mesylate using methanesulfonyl chloride (MsCl), followed by nucleophilic substitution with a bifunctional diazirine-alkyne linker to afford the photoaffinity probe. *(B)* Schematic overview of the affinity-based protein profiling (A*f*BPP) workflow. Intact *D. desulfuricans* cells were treated with 10 μM **DXE**-probe or DMSO as a control, followed by UV-induced photocrosslinking, cell lysis, and CuAAC-mediated conjugation to either TAMRA or biotin. Probe-labeled proteins were analyzed by SDS-PAGE or enriched using streptavidin beads, enzymatically digested, and identified by LC-MS/MS. *(C)* Volcano plot of A*f*BPP experiments comparing **DXE**-probe-treated and DMSO-treated D. desulfuricans cells. The vertical and horizontal dashed lines indicate a log₂ fold enrichment of 1.3 and a −log₁₀(p) value of 1.3, respectively. FliD was identified as the most significantly enriched protein.

### Chemical Proteomics Identifies Flagellar Proteins as Dexamethasone Targets

*D. desulfuricans* cultures were incubated with the *photo*-**DXE** probe (10 µM), followed by UV irradiation to enable covalent crosslinking, cell lysis, and CuAAC with biotin-azide. Labeled proteins were subsequently enriched using streptavidin beads, tryptically digested, and analyzed by liquid chromatography-tandem mass spectrometry (LC-MS/MS) via label-free quantification (Fig. 3*B*). Proteins identified in the analysis were visualized in volcano plots and most prominent candidate hits were selected based on a log_2_ enrichment ratio exceeding 1.3 and a −log₁₀(p) value of 1.3.

Proteomic analysis revealed significant enrichment of several proteins in probe-treated samples compared to DMSO controls (Fig. 3*C*). Notably, the flagellar cap protein FliD (Ddes_0530) emerged as one of the most prominent and consistently recurring hits throughout our analysis. In addition to FliD, proteins associated with membrane-associated processes, cellular metabolism, and redox-related functions were enriched. However, the identification of FliD as a major target is particularly notable, as it provides a direct mechanistic link between **DXE** treatment and the observed defects in bacterial motility.

To complement the findings revealed by A*f*BPP experiments, the whole proteomes of two independent *D. desulfuricans* strains (ATCC 27774, CCUG 72978) were analyzed (Suppl. Fig. S1). Bacteria were treated with 50 µM **DXE** for different times (5, 24, and 48 h). Interestingly, those experiments revealed recurrent alterations in proteins linked to flagellar assembly, motility, and associated membrane and proton motive force (PMF)-dependent processes. While individual proteins were not consistently regulated across all experimental conditions, several flagella-associated factors repeatedly appeared among the significantly altered proteins, including FliS, FliL, FlgG, FlgI, and the flagellar export ATPase FliI, together with proteins involved in PMF generation and cell envelope remodeling. These findings support the hypothesis that **DXE** perturbs the flagellar apparatus and motility-associated pathways in *D. desulfuricans*.

### Structural Modeling Predicts Dexamethasone Binding to FliD

To investigate whether the proteomics-derived FliD target assignment was structurally compatible with ligand binding, we modeled the Ddes_0530-**DXE** complex using AlphaFold3 and Boltz2 (34, 35). AlphaFold3 predicted a hexameric *E. coli* FliD-like assembly for Ddes_0530 (Fig. 4A). To assign the predicted binding site to a functional region, we compared the Ddes_0530 model with an AlphaFold-predicted *E. coli* FliD structure that was annotated according to the domain organization reported by Song et al. for ecFliD (PDB 5H5V) (Fig. 4B, C) (24). Both AlphaFold3 and Boltz2 placed **DXE** in a surface-exposed groove within the D1-associated leg region of Ddes_0530 (Fig. 4D, Suppl. Fig. S2). This region is functionally relevant because the FliD leg was proposed to mimic flagellin D0/D1 elements and to transiently occupy positions subsequently filled by incoming flagellin molecules during filament elongation (24). Ligand engagement at this site could therefore perturb FliD-mediated flagellin incorporation or cap repositioning.

**Figure 4.**
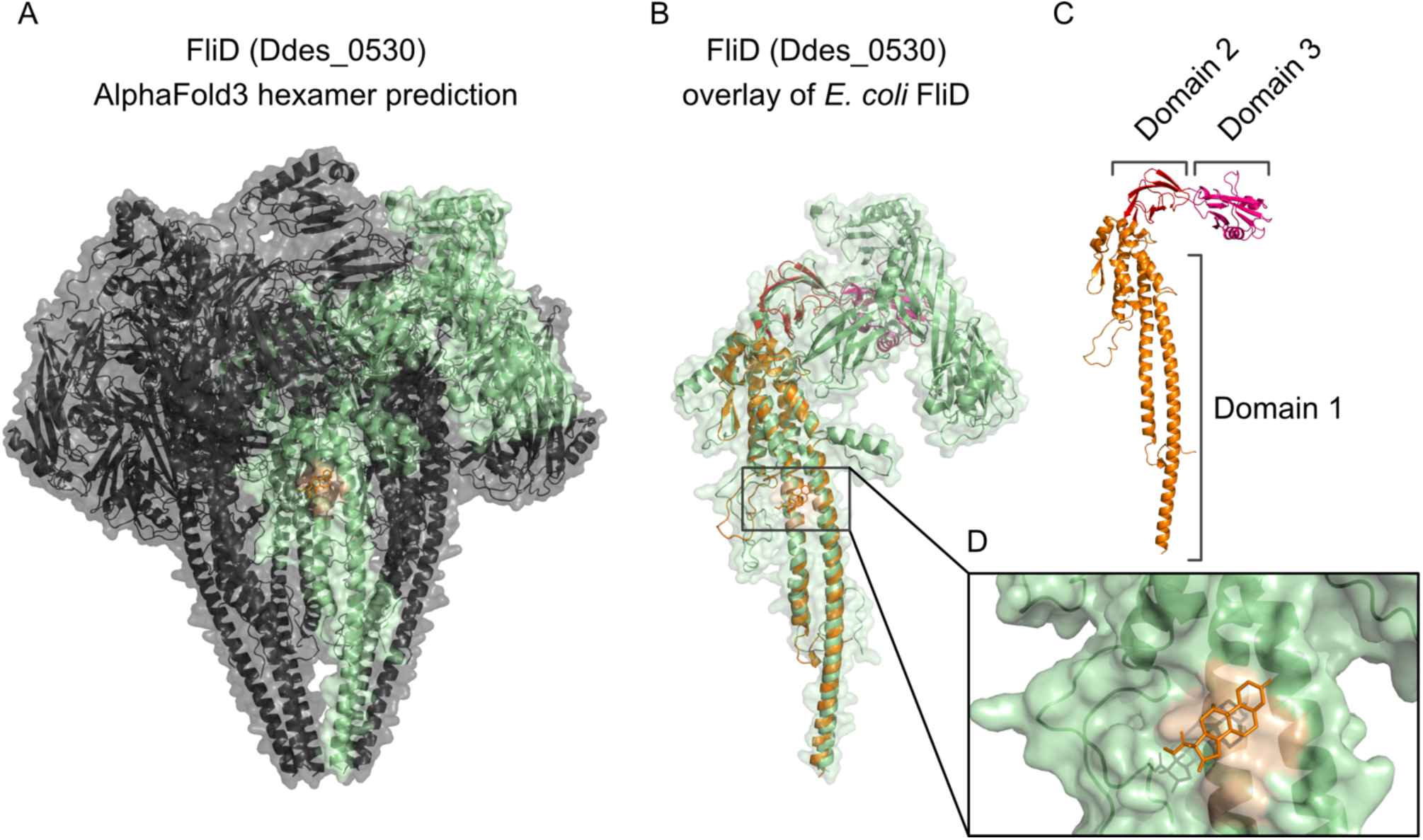
Structural modeling predicts dexamethasone binding to FliD. *(A)* AlphaFold3-predicted hexameric assembly of Ddes_0530, shown as cartoon and transparent surface. One monomer is highlighted in green, while the remaining protomers are shown in gray. The predicted **DXE**-binding site is shown in orange. *(B)* Structural overlay of the AlphaFold3-predicted Ddes_0530 monomer with an AlphaFold-predicted *E. coli* FliD monomer. The Ddes_0530 surface is shown in green and the predicted ligand is shown in orange. *(C)* AlphaFold-predicted *E. coli* FliD monomer annotated according to the domain organization reported for ecFliD by Song et al. (24). The D1 leg domain is shown in orange, while the D2 and D3 cap-plate domains are shown in red and magenta, respectively. *(D)* Close-up of the predicted **DXE**-binding groove in the D1-associated leg region of Ddes_0530. The ligand is shown as orange sticks, and the local binding surface is highlighted.

### Dexamethasone Reduces the Proportion of Flagellated Cells and Flagellar Length

Motile *Desulfovibrio* species possess a single polar flagellum (36). To investigate the functional consequences of **DXE** targeting FliD, we performed transmission electron microscopy (TEM) analysis of flagellar morphology in **DXE**-treated cells (Fig. 5*A*). Flagellar length differed substantially between the two strains under control conditions, with *D. desulfuricans* CCUG 72978 exhibiting longer flagella (6,523 ± 1,041 nm) than *D. desulfuricans* ATCC 27774 (2,971 ± 624 nm; mean ± SD) (Fig. 5*B*). **DXE** reduced the proportion of flagellated cells (Fig. 5*C*). In strain CCUG 72978, the proportion of flagellated cells decreased from 90.3% (control) to 82.3% (**DXE**-treated). A more pronounced effect was observed in strain ATCC 27774, in which the proportion decreased from 83.6% (control) to 71.3% (**DXE**-treated). Treatment with 50 µM **DXE** reduced flagellar length in both strains, reaching statistical significance in ATCC 27774. In **DXE**-treated cells, *D. desulfuricans* CCUG exhibited reduced flagellar length (6,164 ± 1,294 nm), corresponding to a 5.5% decrease compared to controls (Δ = 360 nm; Mann-Whitney test, *p* = 0.066, ns). Similarly, **DXE**-treated strain ATCC 27774 had a mean flagellar length of 2,690 ± 718 nm, representing a 9.4% decrease relative to untreated controls (Δ = 280 nm; Mann-Whitney test, *p* = 0.0006, *).These results strongly support the idea that **DXE** binding to FliD interferes with filament formation (21, 22).

**Figure 5.**
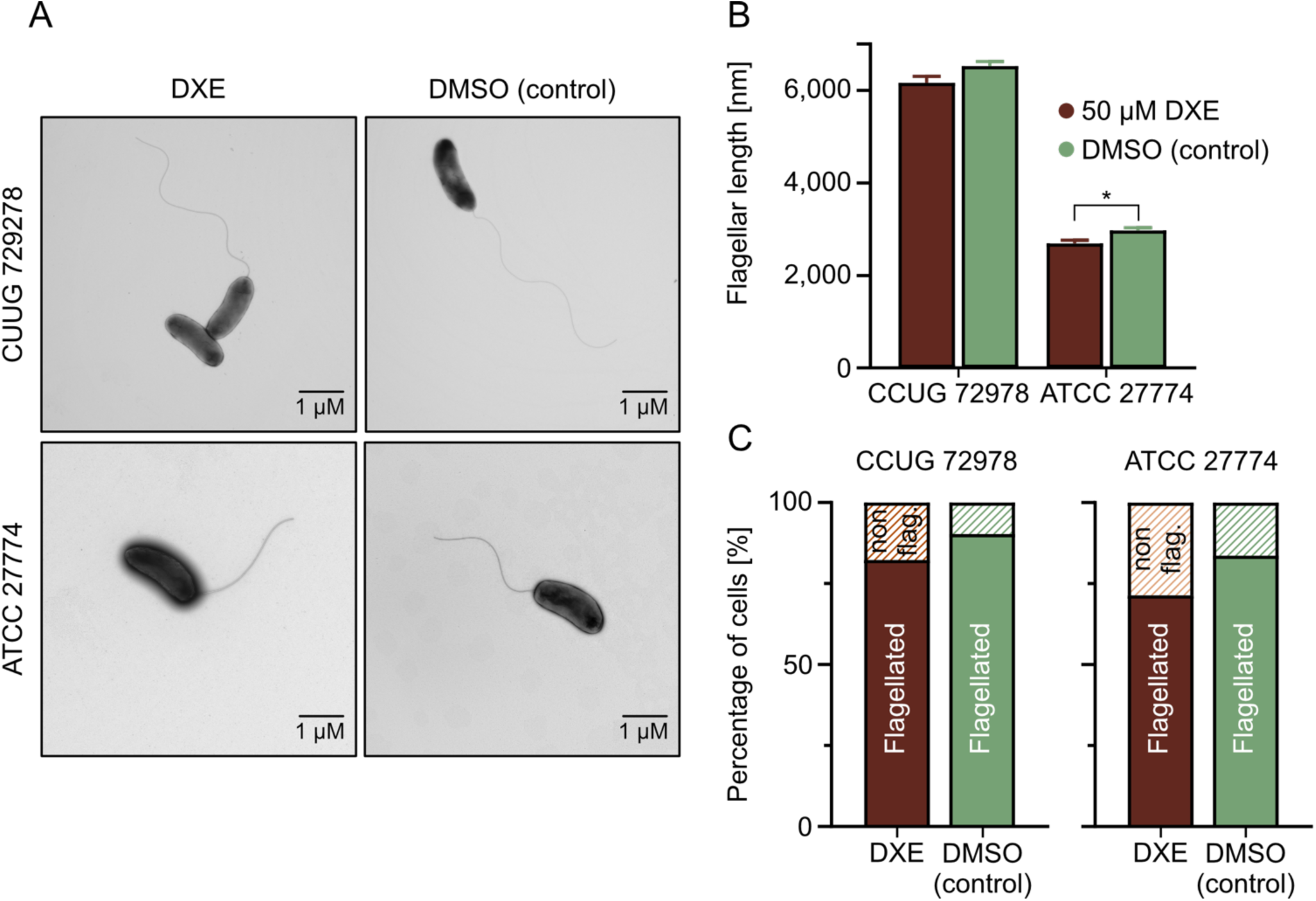
Dexamethasone reduces the proportion of flagellated cells and the length of the flagella. (*A*) Representative transmission electron microscopy (TEM) micrographs of *D. desulfuricans* CCUG 72978 (top) and ATCC 27774 (bottom) grown anaerobically in Postgate medium at 37 °C, either in the presence of 50 µM **DXE** (left) or an equal volume of DMSO for 24 h (right). Cells were negatively stained with 1% (w/v) uranyl acetate and imaged at 80 kV. Scale bars, 1 µm. (*B*) Quantification of flagellar length (nm) measured from TEM micrographs. Each data point represents a single flagellum. For CCUG 72978, flagellar length decreased from 6,523 ± 1041 nm (control; *n* = 95) to 6,164 ± 1294 nm (**DXE**-treated; *n* = 90), a 5.5% reduction (Mann-Whitney test, *p* = 0.066, ns). For ATCC 27774, mean flagellar length decreased from 2,971 ± 624 nm (control; *n* = 120 flagella) to 2,690 ± 718 nm (**DXE**-treated; *n* = 115 flagella), representing a 9.4% reduction (Mann-Whitney test, *p* = 0.0006, *). Error bars represent SEM. (*C*) Proportion of flagellated cells (%) of the indicated strains determined from TEM images.

### Dexamethasone Modulates Swimming Velocity in a Non-Monotonic, Dose-Dependent Manner

To characterize the functional consequences of altered flagellar morphology, we performed light microscopy-based motility tracking of *D. desulfuricans* cells treated with 0, 10 and 50 µM **DXE**. Quantitative single-cell tracking revealed that **DXE** treatment elicited a non-monotonic, dose-dependent effect on swimming velocity (Fig. 6*A*). Control cells (0 µM; n = 1,199) exhibited a mean velocity of 1.93 ± 1.80 µm/s (median 1.53 µm/s). At 10 µM **DXE** (n = 1,970), mean velocity increased 1.9-fold to 3.67 ± 2.56 µm/s (median 3.16 µm/s; *p* < 0.0001, Mann-Whitney *U* test). However, at 50 µM **DXE** (n = 1,037), velocity returned to near-baseline levels (2.17 ± 2.28 µm/s, median 1.44 µm/s; *p* = 0.58 versus control). This biphasic response is consistent with a hormetic mechanism.

**Figure 6.**
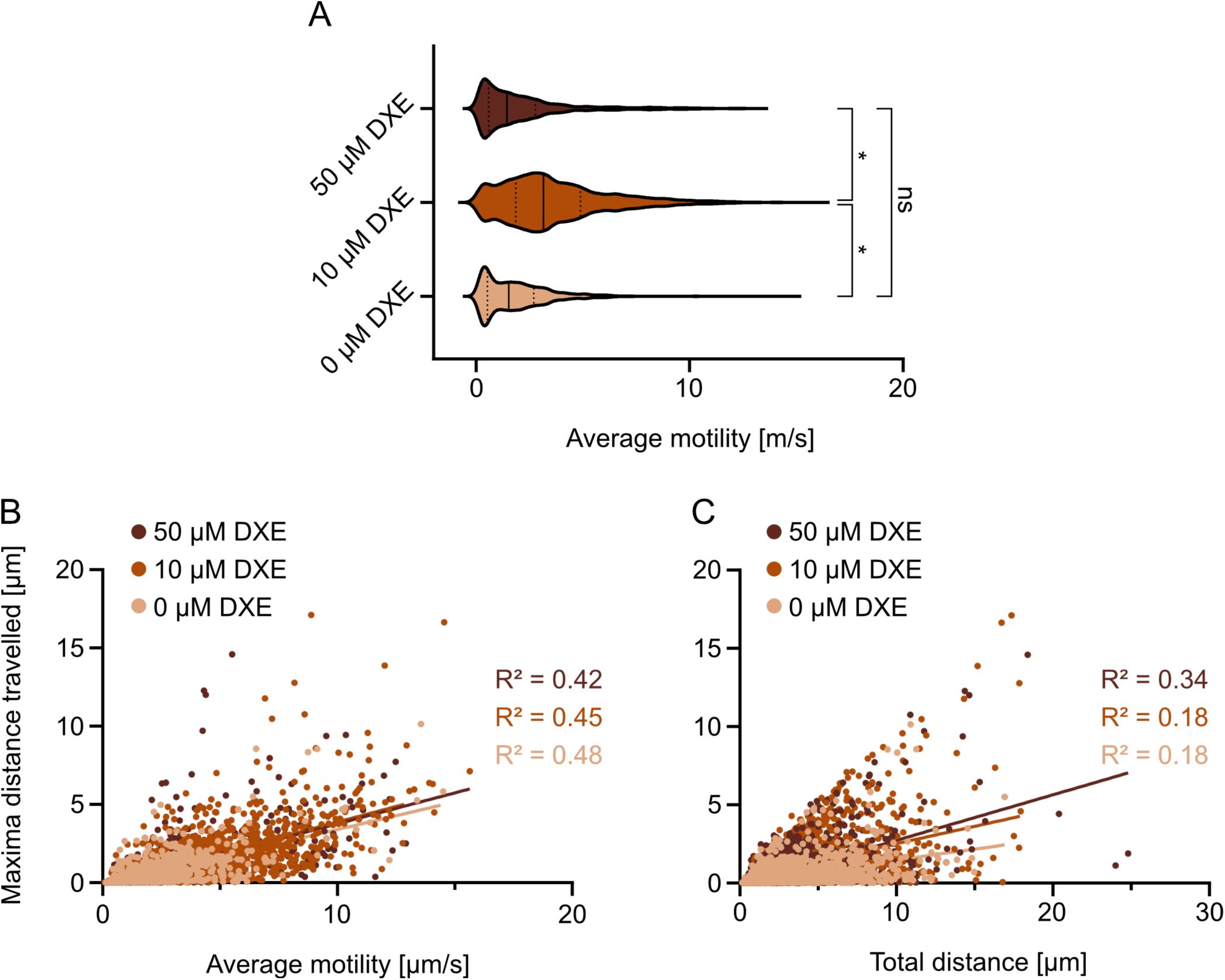
Dexamethasone modulates swimming motility of *D. desulfuricans* in a non-monotonic, dose-dependent manner. (*A*) Violin plots of mean swimming velocity (µm/s) for individual tracked cells across three **DXE** concentrations: 0 µM (control; *n* = 1,199), 10 µM (*n* = 1,970), and 50 µM (*n* = 1,037). Thick bar indicates medians. Treatment with 10 µM **DXE** significantly increased mean velocity (3.67 ± 2.56 µm/s) compared to control (1.93 ± 1.80 µm/s; *p* < 0.0001, Mann-Whitney U test), whereas 50 µM **DXE** returned velocity to near-baseline levels (2.17 ± 2.28 µm/s; *p* = 0.58 versus control). **** *p* < 0.0001; ns, not significant. (*B*) Scatter plot of maximal distance from origin (net displacement, µm) versus mean swimming velocity (µm/s) for each tracked cell, colored by **DXE** condition. Linear regression fits with Pearson correlation coefficient *r*² values are shown for each group (0 µM, *r*² = 0.48; 10 µM, *r*² = 0.45; 50 µM, *r*² = 0.42). The 10 µM **DXE** population (light blue) is shifted toward higher velocities and greater net displacement relative to control (black) and 50 µM (dark blue) populations. (*C*) Scatter plot of maximal distance from origin (net displacement, µm) versus total distance traveled (total path length, µm) for each tracked cell, colored by **DXE** condition. Linear regression fits with Pearson correlation coefficient *r*² values are shown (0 µM, *r*² = 0.18; 10 µM, *r*² = 0.18; 50 µM, *r*² = 0.34). The steeper slope and higher *r*² for the 50 µM condition suggest that, although these cells travel shorter total distances, a larger fraction of their path contributes to net displacement.

The relationship between swimming velocity and spatial displacement further supported the non-monotonic effect of **DXE** on motility (Fig. 6*B*). Scatter plot analysis of net displacement versus mean velocity for each tracked cell revealed a positive linear correlation across all three conditions (Pearson correlation coefficient *r*^2^ = 0.48, 0.45, and 0.42 for 0, 10, and 50 µM **DXE,** respectively). Notably, the 10 µM **DXE** population was shifted toward both higher velocities and greater net displacement relative to control and 50 µM **DXE** populations, indicating that the intermediate dose enhances not only swimming speed but also the capacity for directed spatial exploration. The relationship between net displacement and total path length further supported these findings (Fig. 6*C*). Linear regression analysis revealed *r²* values of 0.18, 0.18 and 0.34 for 0, 10 and 50 µM **DXE** respectively, indicating that at 50 µM a larger fraction of path length contributes to net displacement.

The paradoxical observation of increased swimming velocity despite shorter flagella may reflect altered flagellar mechanics or compensatory changes in motor function. Shorter flagella may be more rigid, enabling more efficient force transmission from the motor to the surrounding medium. The non-monotonic dose-response further suggests that moderate perturbation of FliD function at 10 µM **DXE** may alter flagellar dynamics in a way that favors faster, straighter swimming, whereas higher concentrations (50 µM) may impair flagellar function more severely, returning motility to baseline levels.

### Direct Visualization of Dexamethasone Association with Flagellar Poles

To directly visualize the localization of **DXE** with bacterial cells, we performed fluorescence microscopy using TAMRA-labeled dexamethasone (TAMRA-**DXE**), TAMRA-labeled norethisterone (TAMRA-**NOR**) or TAMRA alone (dye-only control). TAMRA-**NOR** was included as a structurally related steroid control because norethisterone is a commercially available synthetic progestogen that inherently contains an alkyne moiety, allowing direct fluorophore conjugation while enabling us to distinguish **DXE**-specific interactions from nonspecific steroid binding. Live cells were incubated with the different compounds for 10 minutes, washed twice with motility buffer and imaged using phase contrast and Texas Red (TXR) fluorescence microscopy. Fluorescent foci were quantified using the MicrobeJ plugin for ImageJ (37), which detects fluorescent maxima and their spatial relationship to cell poles.

Quantitative analysis revealed a consistent trend across both *D. desulfuricans* strains ATCC 27774 and CCUG 72978. Cells incubated with TAMRA-**DXE** exhibited a higher proportion of polar fluorescent foci compared with TAMRA-only controls (Fig. 7A). In ATCC 27774 and CCUG 72978, the fraction of cells displaying one polar maximum increased after TAMRA-**DXE** treatment, while the fraction with no polar maxima decreased. Given that TEM analysis confirmed that these strains possess a single polar flagellum, the enrichment of TAMRA-**DXE** signal at cell poles is consistent with **DXE** association with the flagellar apparatus.

**Figure 7.**
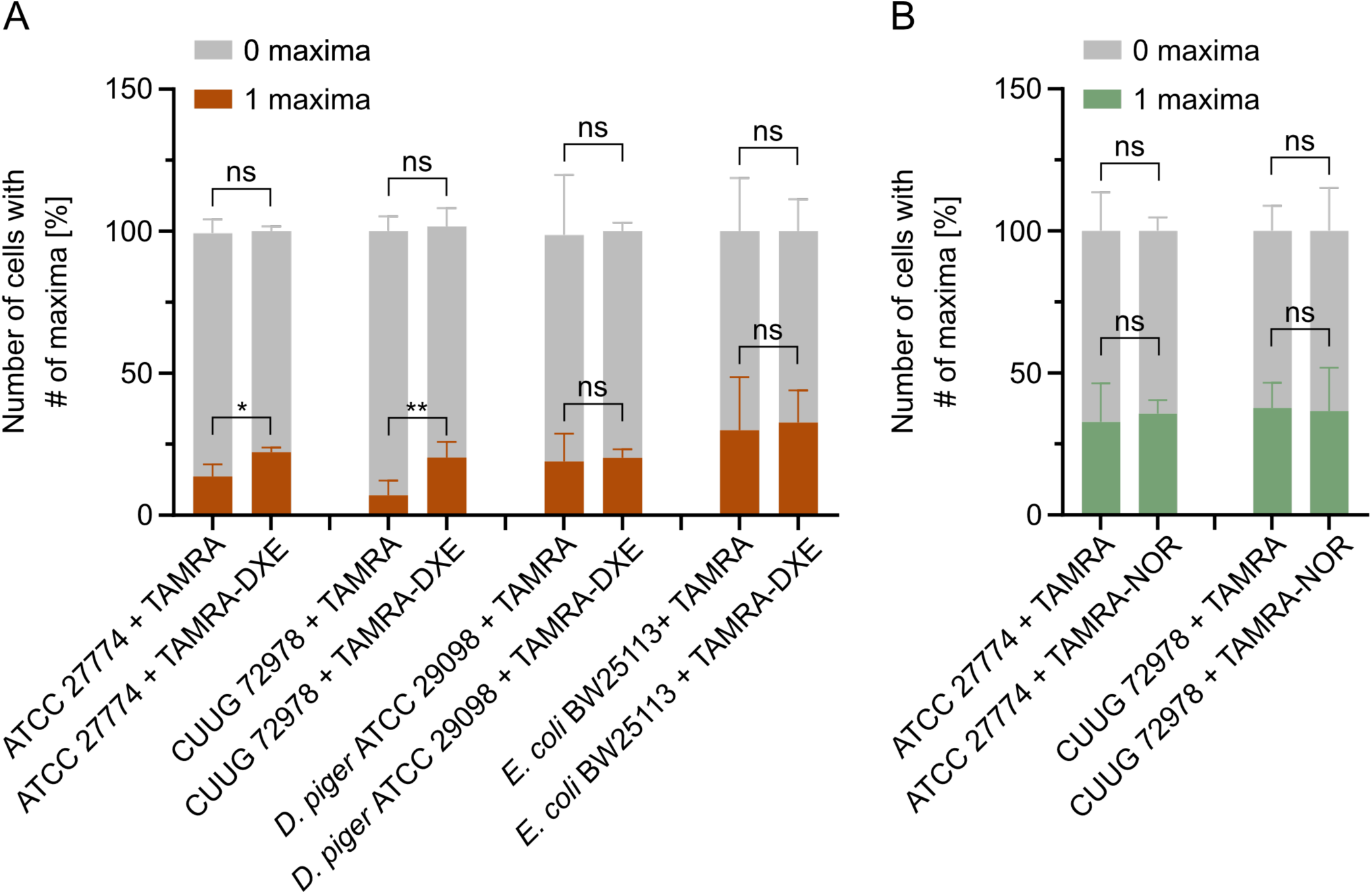
Direct visualization of dexamethasone association with flagellar poles in D. desulfuricans. (*A*) Percentage of cells exhibiting one polar fluorescent focus after incubation with TAMRA-**DXE** or TAMRA in D. desulfuricans ATCC 27774 and CCUG 72978. D. piger ATCC 29098 and E. coli BW25113 wild-type cells served as negative controls. (*B*) Percentage of cells exhibiting one polar fluorescent focus after incubation with TAMRA-NOR (TAMRA-norethisterone) or TAMRA in D. desulfuricans ATCC 27774 and CCUG 72978. Polar fluorescent foci (maxima) were detected automatically and classified according to their spatial relationship to the cell poles using the MicrobeJ plugin in ImageJ/Fiji. At least 1,300 cells were analyzed per strain and condition. Statistical analysis was performed using a two-tailed unpaired t-test. \**p* = 0.0326; \*\**p* = 0.0387; ns, not significant.

Importantly, this polar localization pattern was not observed in *D. piger* (a non-motile, non-flagellated *Desulfovibrio* species) or *E. coli* BW25113 (a flagellated species carrying a FliD homolog) used as controls. Wild-type *D. piger* and *E. coli* BW25113 showed similar proportions of polar fluorescent foci, regardless of whether cells were incubated with TAMRA-**DXE** or TAMRA alone (Fig. 7A). The absence of **DXE**-specific polar localization in *D. piger* and *E. coli* suggests that the observed polar enrichment in *D. desulfuricans* reflects species-specific differences in FliD structure or flagellar accessibility.

Furthermore, a similar steroid compound, norethisterone labeled with TAMRA, did not show a higher proportion of polar fluorescent foci in either of the two *D. desulfuricans* strains compared with TAMRA-only controls (Fig. 7B), indicating the specificity of TAMRA-**DXE** for the flagellar structure of *D. desulfuricans*.

These imaging results provide independent support for a co-localization of **DXE** with the flagellar apparatus in *D. desulfuricans*, complementing our chemical proteomics identification of FliD as a binding target.

## Discussion

In this study, we report the direct targeting of the flagellar cap protein FliD by the synthetic glucocorticoid **DXE** in *D. desulfuricans*. This finding establishes a new paradigm for metabolism-independent drug-microbiome interactions, wherein therapeutic drugs can directly affect bacterial physiology through protein binding rather than metabolic transformation (2, 5).

The identification of FliD as a target of **DXE** was unexpected but provides a mechanistic explanation for the observed effects on flagellar morphology and motility. FliD is essential for flagellar assembly, forming oligomeric caps at the growing filament tip that facilitate flagellin polymerization (22, 24). Our structural modeling predicts that **DXE** binds to a groove in the C-terminal domain of FliD, a region that extends from the core structure and may be involved in oligomerization. Dysfunction of FliD would impair flagellar assembly, consistent with our observations of reduced flagellar length and an increase of the number of non-flagellated cells.

The metabolic stability of **DXE** in *Desulfovibrio* cultures distinguishes this organism from steroid-metabolizing bacteria such as *C. steroidoreducens*, which encodes the OsrABC pathway capable of reducing synthetic glucocorticoids to inactive metabolites (6, 38). The absence of detectable **DXE** metabolism in *Desulfovibrio* indicates that the observed effects result from direct drug-protein interaction rather than metabolic transformation. This has important implications for understanding drug-microbiome interactions, as it suggests that the effects of glucocorticoids on gut bacteria may extend beyond the subset of organisms capable of steroid metabolism (39, 40).

The paradoxical observation of increased swimming velocity despite shorter flagella warrants further investigation. Notably, the motility response was non-monotonic: the 10 µM **DXE** condition produced the strongest stimulation (1.9-fold increase in mean velocity), while 50 µM **DXE** returned motility to near-baseline levels. One possibility is that, due to FliD perturbation, the flagella are not only shorter, but also change in stiffness and/or motor-filament coupling in a way that favors faster, straighter swimming, whereas more severe disruption at higher concentrations impairs propulsion (41). The straighter swimming paths observed in **DXE**-treated cells could result from changes in the flagellar switching frequency or motor torque (42, 43).

*Desulfovibrio* species are sulfate-reducing bacteria increasingly recognized as key players in gut inflammation, where their flagellin activates the NAIP/NLRC4 inflammasome and the LRRC19 receptor, driving pro-inflammatory cytokine production (12, 44, 45). **DXE**, a glucocorticoid widely used to suppress host inflammation, may paradoxically affect *Desulfovibrio* motility and spatial exploration at therapeutic concentrations (46, 47). Increased motility could potentially enhance bacterial dissemination through the mucus layer and exacerbate flagellin-mediated immune activation, highlighting a complex, dose-dependent off-target effect of glucocorticoids on the gut microbiota (48, 49).

In a broader context, our findings have potential clinical implications for understanding glucocorticoid therapy in IBD patients. *Desulfovibrio* species colonize approximately 50% of human intestines and their abundance has been associated with various gastrointestinal conditions (11, 14, 15). The direct effects of **DXE** on *Desulfovibrio* motility could influence bacterial colonization patterns, but additionally also biofilm formation and interactions with the host epithelium. Furthermore, the strain-specific responses observed in our growth experiments suggest that individual microbiome composition may influence the local effects of glucocorticoid therapy (27, 39).

Our approach, combining chemical proteomics with structural modeling and functional validation, provides a template for investigating direct drug-microbiome interactions (2, 3, 26, 40, 50). As the gut microbiome is increasingly recognized as a modulator of drug efficacy and toxicity, understanding the molecular mechanisms of drug-microbe interactions will be essential for optimizing therapeutic outcomes.

In conclusion, we have identified FliD as a direct target of **DXE** in *D. desulfuricans* and demonstrated that **DXE** treatment disrupts flagellar assembly and alters motility behavior without undergoing bacterial metabolism. These findings reveal a metabolism-independent mechanism of glucocorticoid-microbiome interaction with potential implications for understanding treatment variability in IBD patients.

## Materials and Methods

### Bacterial strains

*D. desulfuricans* ATCC 27774 (51), *D. desulfuricans* CCUG 72978 (unpublished clinical isolate), *D. desulfuricans* DSM 642 (52), *D. piger* ATCC 29098 (53) and *Escherichia coli* BW25113 (54) were used in this study.

### Swim agar assays

5 µL overnight culture of *D. desulfuricans* ATCC 27774 or *D. desulfuricans* CCUG 72978 were dropped into freshly poured swimming motility Columbia agar plates [Casein peptone 12 g/L, meat peptone 5 g/L, yeast extract 3 g/L, beef extract 3 g/L, corn starch 1 g/L, sodium chloride 5 g/L, agar 0.3% (w/v)]. **DXE** was dissolved in DMSO and added as supplement directly to the autoclaved medium before pouring the plate. As a solvent control, the corresponding volume of DMSO was added to plates. Plates were incubated at 37 °C under anaerobic conditions until rings were visible. Pictures were taken with EOS M50 camera (Canon, Tokyo, Japan). For image analysis Image J software was used. Significance was determined by performing a one-way ANOVA with Turkey’s post hoc test.

### *Desulfovibrio* growth in presence of DXE

Individual *Desulfovibrio* cultures were inoculated from an overnight culture and cultivated in an anaerobic chamber (85% N_2_, 5% H_2_, 10% CO_2_). **DXE** and DMSO were added to the corresponding samples at the timepoint of inoculation. Optical density (OD_600_) measurements were first corrected by subtracting the medium background from each well. To enable comparison of growth kinetics across different conditions and starting inoculum densities, a log-baseline normalization was applied to the background corrected data. For each well, the normalized value at time t was calculated as ln(OD(t)) - ln(OD(t_0_)), where OD(t) is the optical density at time t and OD(t_0_) is the optical density at the initial time point (t = 0). This transformation, mathematically equivalent to ln(OD(t)/OD(t_0_)), linearizes exponential growth and ensures all growth curves originate from zero.

### DXE identification through HRMS

Individual *Desulfovibrio* cultures were inoculated from an overnight culture and cultivated in an anaerobic chamber. **DXE** and DMSO were added to the corresponding samples at the timepoint of inoculation. For every timepoint 10 mL of each culture were taken and cells were separated from supernatant by centrifuging the culture 10 min at 5,000 × *g*. Culture supernatants were extracted with ethyl acetate (2 × 8 mL per 10 mL supernatant). The combined organic phase was dried over MgSO_4_ and the solvent was removed under reduced pressure. Dried extracts were resolved in HRMS-grade methanol and filtered through a 0.2 µm PTFE membrane syringe filter (Fisherbrand, USA) prior to LC-HRMS analysis. LC-HRMS measurements were conducted on a Bruker Impact II ultra-high-resolution Q-TOF mass spectrometer with electron-spray ionization (ESI). Water (A) and acetonitrile (B) were used as solvents, both supplemented with 0.1% (v/v) formic acid. Samples were eluted at 0.3 mL/min with a 30 min gradient 5% to 95% B. Bruker Compass DataAnalysis Version 6.1 was used to analyze the data.

### Structural prediction with AlphaFold3 and Boltz2

The full-length amino acid sequence of Ddes_0530 and the chemical structure of **DXE** were used for protein-ligand complex prediction with AlphaFold3 and Boltz2 (34, 35). AlphaFold3 predictions were generated for the Ddes_0530 hexamer in complex with **DXE** using C[C@@H]1C[C@H]2[C@@H]3CCC4=CC(=O)C=C[C@@]4([C@]3([C@H](C[C@@]2([C@]1(C(=O)CO)O)C)O)F)C for the ligand. The top-ranked prediction was selected based on the confidence scores, and ligand placement was assessed from the predicted binding pose and local model confidence. Boltz2 monomer predictions were generated using the same Ddes_0530 sequence and ligand representation. The resulting Boltz2 model was aligned with the AlphaFold3 prediction using PyMOL to assess whether both methods localized **DXE** to the same structural region. For functional annotation of the predicted binding site, the Ddes_0530 model was structurally compared with an AlphaFold-predicted *E. coli* FliD model. The *E. coli* FliD model was annotated according to the D1, D2, and D3 domain organization reported by Song et al. (24), for ecFliD. Structural overlays and figures were generated in PyMOL. The predicted **DXE**-binding site was assigned to the D1-associated leg region based on its position relative to the annotated FliD domain architecture.

### Transmission electron microscopy (TEM)

*Desulfovibrio* strains were cultured in Postgate medium (55) under anaerobic conditions. Cultures were treated with 50 µM **DXE** or the DMSO (equivalent volume as solvent control) for 24 h. 10 mL samples were collected after 10 min by centrifugation at 5,000 × *g*. Samples were resuspended in 900 µL phosphate-buffered saline (PBS, pH 7.0) and chemically fixed with 2.5% (v/v) glutaraldehyde. Copper grids for TEM were coated with a carbon film of 10 nm thickness and additionally hydrophilized. 10 µL bacterial suspension was dropped on the coated side of the grid for 2 min and followed by blotting. For the staining, the carbon-coated side of the grids were washed two times with sterile water and stained for 30 s with 1% (w/v) uranyl acetate. Between each washing and staining step, blotting with filter paper was repeated to remove liquid. Images of the negative stained samples were taken with the transmission electron microscope (EM912, ZEISS microscopy GmbH) at 80 kV.

### Determination of flagellar length

Flagellar length was quantified from transmission electron microscopy images using **FiJi / ImageJ** (version 21.0.7). TIF image files were opened in FiJi and the flagella were traced manually using the *Freehand Line* tool. The traced length was then converted to micrometers using the appropriate pixel-to-distance calibration set from the microscope metadata. For each sample, at least *n = 80* flagella were measured, and the average length and standard deviation were calculated.

### Microscopic motility tracking

A pre-culture of *D. desulfuricans* ATCC 27774 was inoculated and cultivated in the presence of the given concentrations of **DXE** for 4 h. The culture was transferred to a microscopy slide, and covered with a coverslip (LifterSlip™, EMS, catalog no. E72186-36). The bacterial motility was tracked at 37 °C for 10 s (8.7 fps) on a Leica DMi8 inverted microscope equipped with a Leica DFC365 FX camera (Wetzlar, Germany) and 40× objective lens. Per slide, at least three different positions were tracked. The measurement was performed in independent replicates. For further processing, a custom Python script was used (code available at https://github.com/MIsselstein/E.Coli_Tracker (56)). First, the pictures were defined as binary images and the 99 pictures of a 10 s tracking experiment were put into one movie. For cell detection, the search range (defined as the maximum distance features that can move between frames) was set to 25 and the memory (defined as the maximum number of frames during which a feature can vanish, then reappear nearby, and be considered the same particle) to 2. The magnification was set to 40 and the pixel size to 6.45. The detected trajectories were exported as a movie and as an Excel file for further processing.

### TAMRA-labeled fluorescence microscopy

*D. desulfuricans* ATCC 27774 and CCUG 72978, *D. piger* ATCC 29098, and *Escherichia coli* BW25113 were used for staining experiments with TAMRA-labeled dexamethasone (TAMRA-**DXE**), TAMRA-labeled norethisterone (TAMRA-NOR), or TAMRA dye alone. TAMRA-**DXE**, TAMRA-NOR, and TAMRA were diluted to 300 μM in motility buffer (10 mM potassium phosphate, pH 7.0, 67 mM NaCl, 0.1 mM EDTA). *D. desulfuricans* strains were grown anaerobically in 10 mL Postgate medium for 24 h, whereas *D. piger* was grown in Desulfovibrio (MV) medium (57) for 48 h. *E. coli* was grown in LB medium to an OD_600_ of 0.8. Cultures were harvested by centrifugation at 1,400 × *g* for 8 min at room temperature, and the pellets were resuspended in 100 μL motility buffer. For each strain, 20 μL aliquots were transferred to microcentrifuge tubes and incubated with 50 μM TAMRA-**DXE**, 50 μM TAMRA-NOR, or an equivalent volume of TAMRA dye as a control. After incubation for 10 min at room temperature, samples were washed twice with 1 mL motility buffer and resuspended in 20 μL motility buffer. For microscopy, 10 μL of each suspension was placed on a microscope slide, covered with a coverslip, and allowed to settle for 30 min before imaging. Cells were then examined by phase-contrast and fluorescence microscopy using a TXR filter cube. For TAMRA fluorescence, an excitation wavelength of 546 nm and a 605 nm emission filter with a 75-nm bandwidth were used with an exposure of 100 ms, a gain of 1, and 100% intensity. To quantify polar fluorescent foci (maxima) per cell pole representing TAMRA-**DXE**, TAMRA-NOR, or TAMRA of single cells, phase contrast and fluorescent images were analyzed using the ImageJ (58) plugin MicrobeJ (37). Default settings of MicrobeJ were used for cell segmentation (fit shape, rod-shaped bacteria), except for the following settings: area: 0.1-max µm^2^; length: 1.2-20 µm; width: 0-1.6 µm; and angularity: 0.0-0.7. The number and position/distance to the pole of fluorescent foci (maxima) corresponding to TAMRA-**DXE**, TAMRA-NOR, or TAMRA were determined using the default settings of MicrobeJ “maxima detection” (Foci, Smoothed), except for the following settings: tolerance: 50; z-score: 3; area: 0.1-0.3; intensity: 250-max; outside: “Dist.” 3 µm or 7 µm boundary. The distance threshold used to assign a fluorescent focus (maxima) to a cell pole was 7 µm for *D. desulfuricans* CCUG 72978 and *E. coli* BW25113, and 3 µm for the other tested strains. The flagellar length used for these thresholds in *D. desulfuricans* ATCC 27774 and CCUG 72978 was based on flagellar length measurements obtained from TEM micrographs (Fig. 5B), and for *E. coli* based on published literature values (59). At least 1,300 cells were analyzed per strain and condition.

### Labeling, Cell Lysis, and Click Chemistry

For each biological replicate, *Desulfovibrio desulfuricans* cells were cultured anaerobically in Postgate medium supplemented with 1.8 mM FeSO₄ for 24 h. The resulting preculture was used to inoculate four independent 40 mL cultures of fresh Postgate medium lacking FeSO₄, followed by incubation for an additional 24 h. Due to the formation of iron sulfide precipitates during growth, optical density measurements were not feasible. The culture was harvested by centrifugation at 6,000 × *g* for 10 min at 4 °C, and the resulting pellets were resuspended in 10 mL Postgate medium lacking FeSO₄.

For affinity-based protein profiling (A*f*BPP) experiments, 200 µL aliquots of the cell suspension were transferred into Protein LoBind tubes (Eppendorf) and treated with *photo*-**DXE** (final concentration 10 µM) or DMSO as a control using a final DMSO concentration of 1%. Samples were incubated for 10 min at 37 °C and 200 rpm under aerobic conditions, followed by UV irradiation (FL8BL-B lamps, *Hitachi*) for 5 min under cooling conditions for photo-crosslinking. Cells were subsequently harvested by centrifugation at 6,000 × *g* for 10 min at 4 °C. Cell lysis was performed in 200 µL PBS containing 1% Triton X-100 by probe sonication (3 × 15 s at 80% intensity; Sonopuls HD 2070 ultrasonic rod, *Bandelin electronic GmbH*). Lysates were clarified by centrifugation at 21,300 × *g* for at least 30 min at 4 °C, and the supernatants were transferred into fresh Protein LoBind tubes. Protein concentrations were determined using a bicinchoninic acid (BCA) assay (Roti Quant, *Roth*) according to the manufacturer’s instructions. Protein samples were adjusted to 2.23 µg/µL, and 45 µL of each sample (∼100 µg total protein) was transferred into a 96-well polypropylene V-bottom plate (*Greiner*, cat. 651201).

A click chemistry master mix was prepared by combining 0.6 µL of 20 mM biotin-azide in DMSO, 2.5 µL of 1.67 mM tris(benzyltriazolylmethyl)amine (TBTA) in 80% tert-butanol and 20% DMSO, 1.2 µL of 50 mM copper(II) sulfate (CuSO₄) in water, and 0.6 µL of 100 mM tris(2-carboxyethyl)phosphine (TCEP) in water. Subsequently, 4.9 µL of the click master mix was added to each protein sample, followed by incubation for 90 min at room temperature with shaking at 950 rpm. The reaction was quenched by addition of 65 µL of 8 M urea containing 10 mM TCEP and 20 mM iodoacetamide (IAA), and samples were incubated for 15 min at room temperature and 950 rpm. Residual IAA was subsequently quenched by addition of 2 µL of 500 mM dithiothreitol (DTT) per sample.

### Enrichment, Digestion, and LC-MS/MS Analysis

Protein sample cleanup and enrichment were performed using an adapted single-pot solid-phase-enhanced sample preparation (SP2E) workflow employing carboxylate-coated magnetic beads (60). To each sample, 10 µL of a 1:1 mixture of hydrophobic and hydrophilic carboxylate-coated magnetic beads (*Cytiva*, cat. 65152105050250 and 45152105050250), pre-washed three times with water, was added, followed by precipitation of proteins by addition of 175 µL 100% ethanol. Samples were subsequently processed on an automated liquid handling platform (Hamilton Microlab Prep, *Hamilton*). Beads were washed three times with 180 µL of 80% ethanol and once with 180 µL acetonitrile prior to protein elution in 75 µL of 0.2% SDS in PBS. The elution step was repeated once, resulting in a total elution volume of 150 µL. Biotinylated proteins were enriched using 50 µL streptavidin magnetic beads (*New England Biolabs*, cat. S1420S) pre-washed three times with 0.2% SDS in PBS. Samples were incubated for 1 h at room temperature and 800 rpm to allow binding of labeled proteins to the beads. Subsequently, beads were washed three times with 180 µL of 0.1% NP-40 in PBS, two times with 180 µL of 6 M urea, and three times with 200 µL water.

On-bead digestion was carried out overnight at 37 °C and 800 rpm in 100 µL of 50 mM triethylammonium bicarbonate (TEAB) using sequencing-grade trypsin (*Promega*) at an enzyme-to-protein ratio of 1:100. Peptides were subsequently eluted using 50 µL of 3% formic acid. Desalting was performed using styrenedivinylbenzene-reverse phase sulfonate (SDB-RPS) stage tips (Empore, *3M*) as previously described (61). Eluted peptides were dried using a centrifugal evaporator (Concentrator Plus, Eppendorf) and reconstituted in 35 µL of 1% formic acid prior to LC-MS/MS analysis. A total of 4 µL of each sample was injected for LC-MS/MS measurements. Samples were analyzed on a timsTOF Pro mass spectrometer (*Bruker*) coupled to an UltiMate 3000 UHPLC system (*Thermo Fisher Scientific*) in data-independent acquisition (DIA) mode. All preparative MS experiments were conducted in four biologically independent replicates.

### Full Proteome Analysis

*Desulfovibrio desulfuricans* was cultured anaerobically in Postgate medium supplemented with 1.8 mM FeSO₄ for 24 h. The resulting preculture was used to inoculate four independent 10 mL cultures of fresh Postgate medium lacking FeSO₄, followed by incubation for an additional 24 h. For treatment experiments, cultures were incubated with either **DXE** (50 µM final concentration) or DMSO as a control (1% final DMSO concentration) for 5 h, 24 h, or 48 h. Subsequently, each culture was harvested by centrifugation at 6,000 × *g* for 10 min at 4 °C.

Cell pellets were resuspended in 200 µL PBS containing 1% Triton X-100 and lysed by probe sonication (3 × 15 s at 80% intensity; Sonopuls HD 2070 ultrasonic rod, *Bandelin electronic GmbH*). Lysates were clarified by centrifugation at 21,000 × *g* for at least 30 min at 4 °C, and the supernatants were transferred into fresh Protein LoBind tubes. Protein concentrations were determined using a bicinchoninic acid (BCA) assay (Roti Quant, *Roth*). Protein samples were adjusted to a total protein amount of 20 µg in 80 µL PBS and transferred into a 96-well polypropylene V-bottom plate (*Greiner*, cat. 651201). Reduction and alkylation were performed by addition of 3 µL of a 1:2 mixture of 500 mM tris(2-carboxyethyl)phosphine (TCEP) and 500 mM iodoacetamide (IAA), followed by incubation for 15 min at room temperature and 950 rpm. Residual IAA was subsequently quenched by addition of 2 µL of 500 mM dithiothreitol (DTT) per sample.

Proteins were precipitated onto 10 µL of a 1:1 mixture of hydrophobic and hydrophilic carboxylate-coated magnetic beads (*Cytiva*, cat. 65152105050250 and 45152105050250), pre-washed three times with water, by addition of 150 µL ethanol and processed on an automated liquid handling platform (Hamilton Microlab Prep, *Hamilton*). Beads were washed three times with 180 µL of 80% ethanol and once with 180 µL acetonitrile prior to overnight on-bead digestion in 100 µL of 50 mM triethylammonium bicarbonate (TEAB) using sequencing-grade trypsin (*Promega*) at an enzyme-to-protein ratio of 1:100 at 37 °C and 800 rpm. Peptides were eluted using 50 µL 3% formic acid and desalted using SDB-RPS stage tips (Empore, *3M*) as previously described (61). Eluted peptides were dried using a centrifugal evaporator (Concentrator Plus, *Eppendorf*) and reconstituted in 50 µL of 1% (v/v) formic acid prior to LC-MS/MS analysis. Samples were analyzed on a timsTOF Pro mass spectrometer (*Bruker*) coupled to an UltiMate 3000 UHPLC system (*Thermo Fisher Scientific*) in data-independent acquisition (DIA) mode. A total of 4 µL of each sample was injected for LC-MS/MS measurements.

### LC-MS/MS Measurements and Data Analysis with timsTOF Pro

#### LC-MS/MS measurements on timsTOF Pro

Peptides were processed and separated using an UltiMate 3000 nano HPLC system (*Dionex*) linked to a *Bruker* timsTOF Pro mass spectrometer via a CaptiveSpray nano-electrospray ion source and a *Sonation* column oven. Initially, peptides were loaded onto a trap column (Acclaim PepMap 100 C18, 75 µm ID × 2 cm, 3 µm particle size, *Thermo Fisher Scientific*) and rinsed with 0.1% formic acid in H_2_O for 7 min at a flow rate of 5 µL/min. They were then transferred to a separation column (Aurora C18 column, 25 cm × 75 µm, 1.7 µm, *IonOpticks*) and separated with a gradient: 5% to 17% B over 36 min, 17% to 25% B over 18 min, 25% to 37% B over 6 min, followed by 10 min at 95% B before re-equilibration, at a flow rate of 400 nL/min. Mobile phase A was 0.1% formic acid in H_2_O, and mobile phase B was 0.1% formic acid in acetonitrile. The timsTOF Pro was configured to operate in data-independent dia-PASEF mode, utilizing a dual TIMS analyzer with equal accumulation and ramp times of 100 ms each. The 1/K0 ion mobility ranged from 0.60 to 1.60 V × s/cm² for MS1 scans. Fragmentation was accomplished with the following dia-PASEF settings: a mass range of 400 to 1,201 m/z and an ion mobility range of 0.60 to 1.43 V × s/cm². Ion mobility isolation windows for each dia-PASEF scan were of 26 m/z widths. By employing 32 isolation windows with 1 m/z overlaps, the setup spanned the entire mass range and resulted in 16 dia-PASEF scans per MS1 scan. The total cycle time was about 1.8 s (Table 1). The collision energy was linearly ramped from 59 eV at 1/K0 = 1.3 V × s/cm² to 20 eV at 1/K0 = 0.85 V × s/cm². The TIMS elution voltages were linearly calibrated with an Agilent ESI-L Tuning Mix (ions with m/z of 622, 922, and 1,222) added to the CaptiveSpray Source inlet filter to get the reduced ion mobility coefficients (1/K0).

**Table 1:**
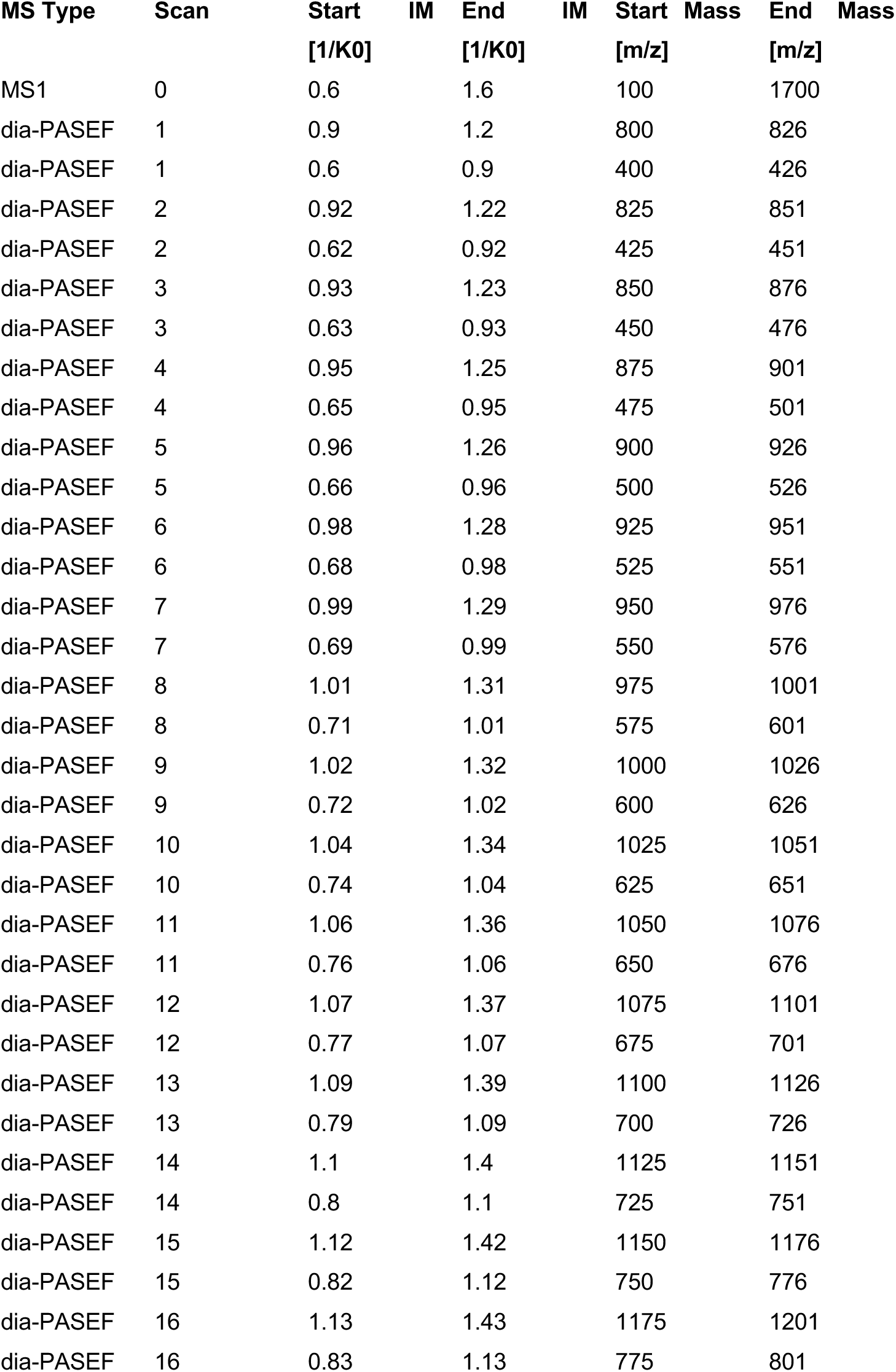
Overview about DIA-PASEF scan windows with ion mobility range (1/K_0_) and scan width (m/z).

#### Data analysis of timsTOF Pro measurements

MS data were processed with DIA-NN (62) (version 1.8.1) using a library-free mode and the UniProt reference proteome for *D. desulfuricans* DSM6949 (taxon identifier: 525146, downloaded on 25.12.2022) was employed to generate the library. Precursor ion settings were the following: library creation and the use of deep-learning algorithms to predict spectra, retention times (RTs), and ion mobilities (IMs). Trypsin/P was selected as the protease, allowing for up to two missed cleavages. The method included N-terminal methionine excision and fixed carbamidomethylation of cysteines, with no variable modifications applied. Peptides ranged from 7 to 30 residues in length, and precursor charges were set between 2 and 4. The precursor m/z range was established from 300 to 1,800, and the fragment m/z range was from 200 to 1,800 for TIMS data. The precursor false discovery rate (FDR) was maintained at 0.01. Mass accuracy, MS1 accuracy, and scan window settings were all configured to 0. Features like isotopologues, match-between-runs (MBR), and removal of potential interferences were enabled. The neural network classifier worked in single-pass mode, performing protein inference at the gene level with heuristic protein inference activated (--relaxed-prot-inf). Quantification utilized the robust LC (high precision) strategy. Cross-run normalization depended on RT, smart profiling was used for library generation, and settings were optimized for both speed and RAM usage. Following DIA-NN analysis, LFQ quantities for all protein groups were examined using Perseus software (63) (version 2.03.1). LFQ intensities were transformed to log_2_ values, and protein groups were filtered to include those with at least three valid values in at least one group. The data was imputed from a normal distribution with default settings (width = 0.3 and down shift = 1.8 for total matrix). A two-sample Student’s t-test was conducted for all relevant comparisons, and fold change values and statistical significance were determined. Graphs were generated using GraphPad Prism 10.01.

## Supplementary Information

### Synthesis

**2-((8S,9R,10S,11S,13S,14S,16R,17R)-9-fluoro-11,17-dihydroxy-10,13,16-trimethyl-3-oxo-6,7,8,9,10,11,12,13,14,15,16,17-dodecahydro-3H-cyclopenta[a]phenanthren-17-yl)-2-oxoethyl methanesulfonate**

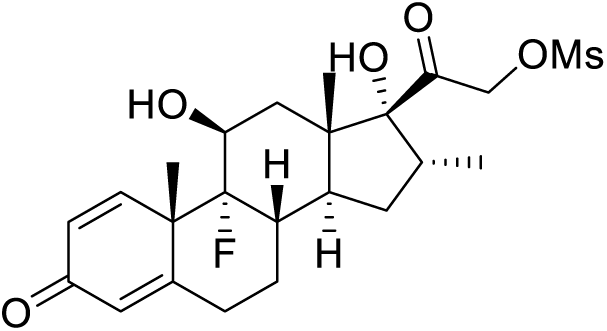

To a solution of dexamethasone (294 mg, 0.75 mmol) in 5 mL anhydrous pyridine MsCl (0.697 mL, 9 mmol) was added dropwise at 0 °C, then the reaction was stirred at 0 °C for 5 h. The reaction was quenched with 5 mL water, extracted with ethyl acetate (3 × 10 mL), washed with 20 mL saturated brine, and concentrated in vacuo. The residue mixture was purified with silica gel chromatography (pentane: ethyl acetate = 1: 1) to afford the 2-((8S,9R,10S,11S,13S,14S,16R,17R)-9-fluoro-11,17-dihydroxy-10,13,16-trimethyl-3-oxo-6,7,8,9,10,11,12,13,14,15,16,17-dodecahydro-3H-cyclopenta[a]phenanthren-17-yl)-2-oxoethyl methanesulfonate (315 mg, 89%) as a white solid

**^1^H NMR (400 MHz, DMSO)** δ 7.29 (d, *J* = 10.1 Hz, 1H), 6.22 (dd, *J* = 10.1, 1.8 Hz, 1H), 6.01 (t, *J* = 1.8 Hz, 1H), 5.38 (dd, *J* = 4.7, 1.6 Hz, 1H), 5.29 (d, *J* = 17.8 Hz, 1H), 5.27 (s, 1H), 4.90 (d, *J* = 17.8 Hz, 1H), 4.19 – 4.09 (m, 1H), 3.24 (s, 3H), 2.90 (tq, *J* = 10.8, 6.7, 5.4 Hz, 1H), 2.62 (td, *J* = 13.6, 6.0 Hz, 1H), 2.44 – 2.27 (m, 2H), 2.20 – 2.06 (m, 2H), 1.83 – 1.73 (m, 1H), 1.65 (q, *J* = 11.7 Hz, 1H), 1.48 (s, 3H), 1.46 (d, *J* = 10.8 Hz, 1H), 1.35 (qd, *J* = 12.7, 4.9 Hz, 1H), 1.08 (ddd, *J* = 12.0, 8.0, 4.0 Hz, 1H), 0.90 (s, 3H), 0.81 (d, *J* = 7.2 Hz, 3H).

**^13^C NMR (101 MHz, DMSO)** δ 203.68, 185.29, 167.03, 152.73, 129.03, 124.13, 101.21 (d, *J* = 175.3 Hz), 90.60, 72.80, 70.49 (d, *J* = 36.8 Hz), 47.99, 47.93 (d, *J* = 22.9 Hz), 43.32, 37.84, 35.73, 35.46, 33.59 (d, *J* = 19.2 Hz), 31.90, 30.26, 27.28, 22.95 (d, *J* = 5.6 Hz), 16.52, 15.10.

**^19^F NMR (376 MHz, DMSO)** δ-164.48.

**HRMS (ESI)** m/z: [M+1]^+^ calc. 471.1848, found 471.1845

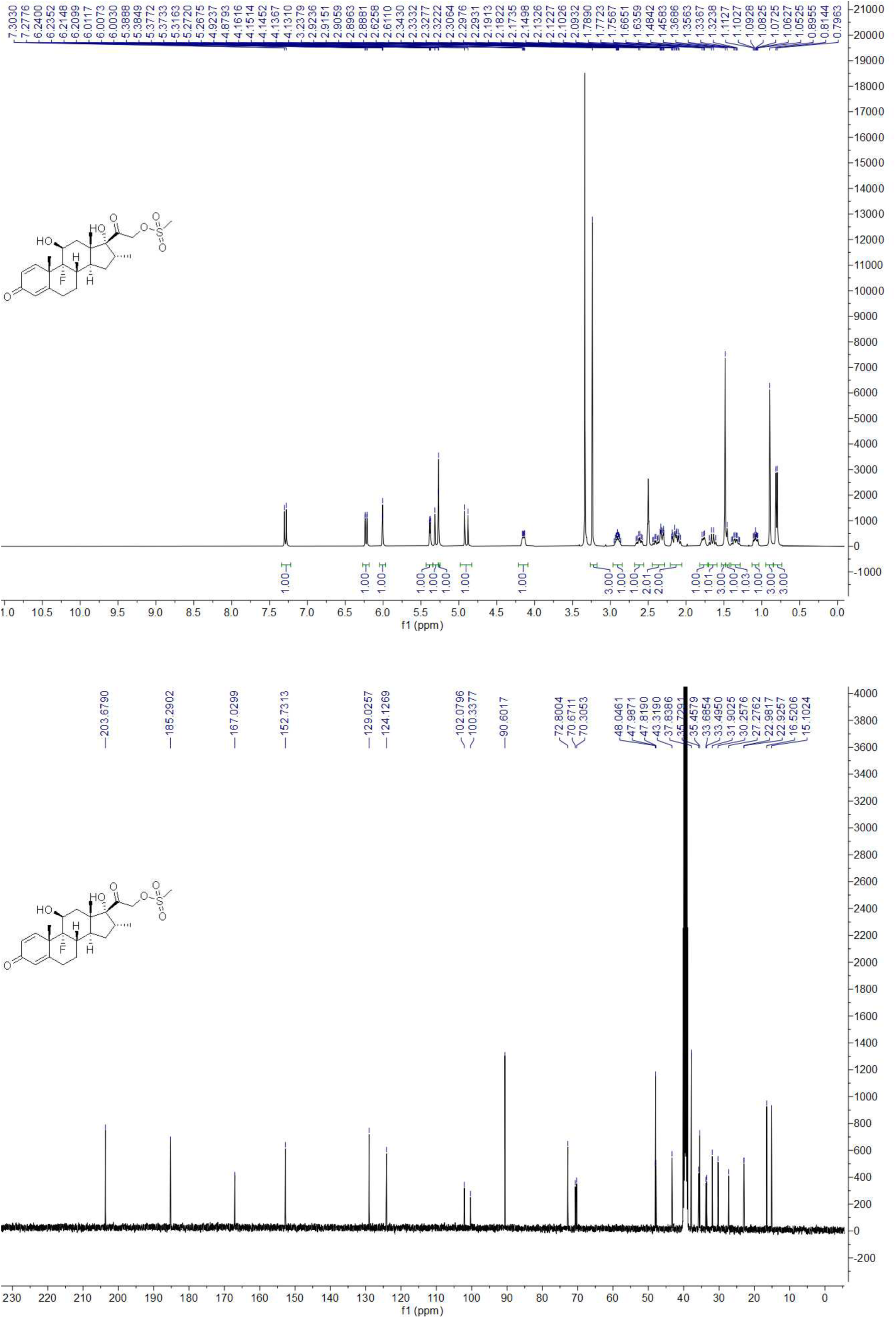

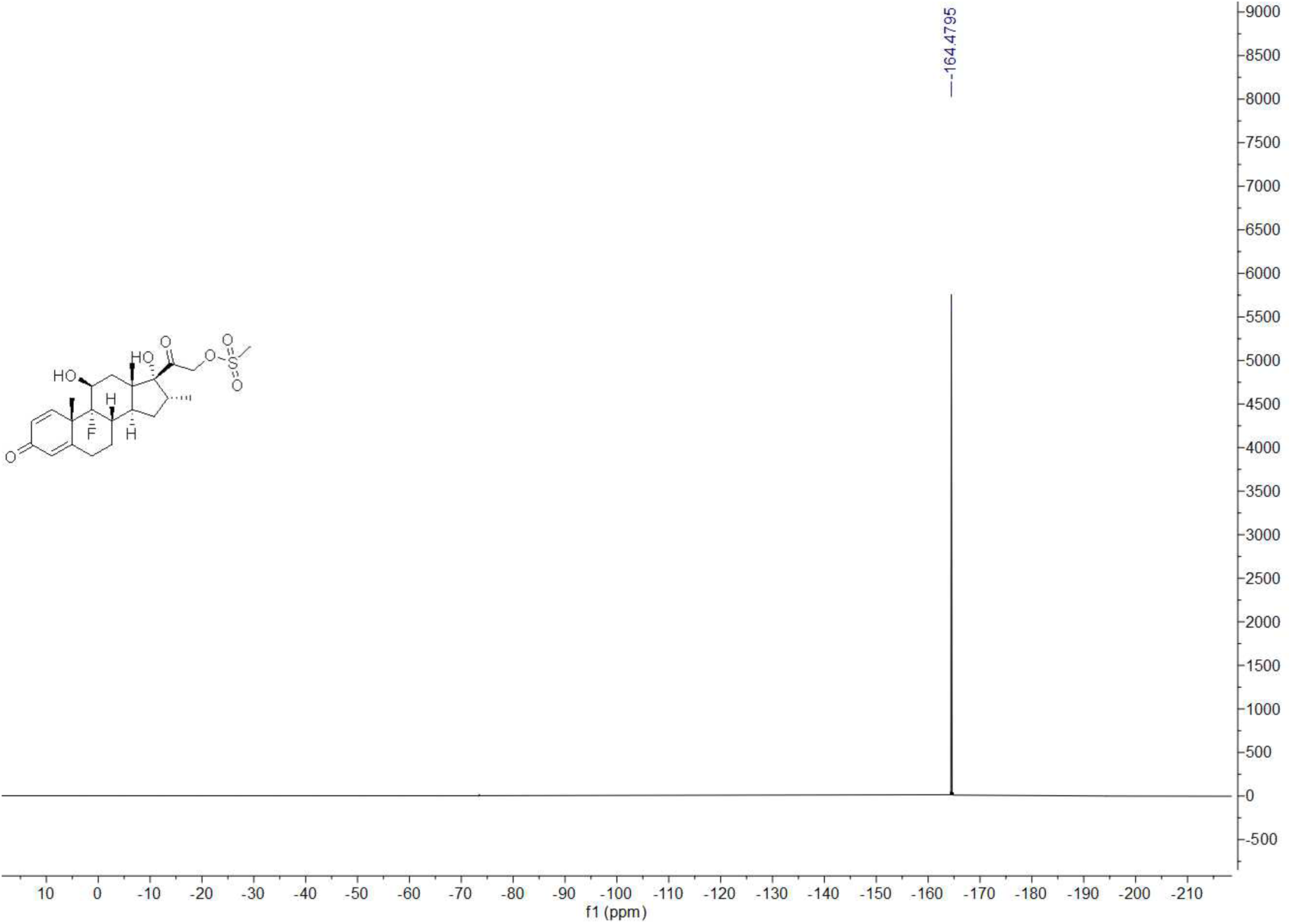

**N-(2-(3-(but-3-yn-1-yl)-3H-diazirin-3-yl)ethyl)-2-((8S,9R,10S,11S,13S,14S,16R,17R)-9-fluoro-11,17-dihydroxy-10,13,16-trimethyl-3-oxo-6,7,8,9,10,11,12,13,14,15,16,17-dodecahydro-3H-cyclopenta[a]phenanthren-17-yl)-2-oxoethan-1-aminium 2,2,2-trifluoroacetate**

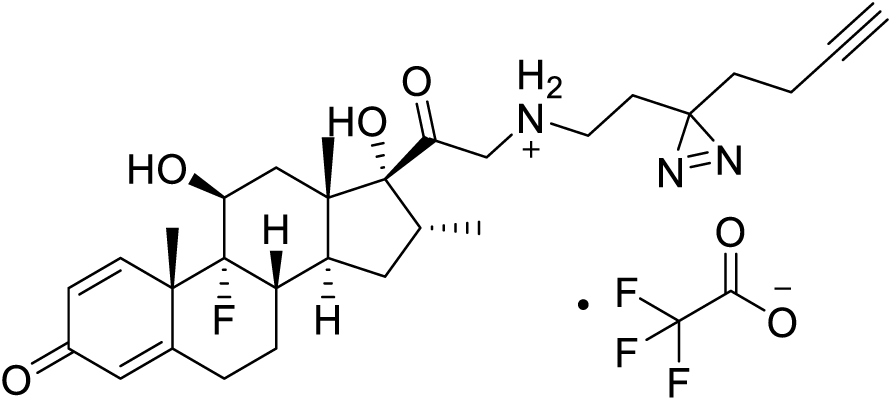

To a solution of 2-((8S,9R,10S,11S,13S,14S,16R,17R)-9-fluoro-11,17-dihydroxy-10,13,16-trimethyl-3-oxo-6,7,8,9,10,11,12,13,14,15,16,17-dodecahydro-3H-cyclopenta[a]phenanthren-17-yl)-2-oxoethyl methanesulfonate (70 mg, 0.15 mmol) and DIPEA (52 µL, 0.3 mmol) in 1.0 mL anhydrous DMF was added 2-(3-(but-3-yn-1-yl)-3H-diazirin-3-yl)ethan-1-amine (31 mg, 0.225 mmol) at room temperature under argon atmosphere, then the reaction was stirred at 65 °C for 8 h under argon atmosphere. The reaction was cooled down to room temperature, quenched with 5 mL water, extracted with ethyl acetate (3 ×10 mL), washed with 20 mL sat. brine, concentrated in vacuo. The reaction residue was purified with preparative HPLC (C_18_, from 2: 98 to 40: 60, 0.1% TFA in CH_3_CN: 0.1% TFA in H_2_O) to afford the N-(2-(3-(but-3-yn-1-yl)-3H-diazirin-3-yl)ethyl)-2-((8S,9R,10S,11S,13S,14S,16R,17R)-9-fluoro-11,17-dihydroxy-10,13,16-trimethyl-3-oxo-6,7,8,9,10,11,12,13,14,15,16,17-dodecahydro-3H-cyclopenta[a]phenanthren-17-yl)-2-oxoethan-1-aminium 2,2,2-trifluoroacetate (25 mg, 26%) as a white solid.

**^1^H NMR (500 MHz, CD_3_CN)** δ 7.24 (d, *J* = 10.1 Hz, 1H), 6.22 (dd, *J* = 10.1, 1.9 Hz, 1H), 6.00 (t, *J* = 1.9 Hz, 1H), 4.26 (dq, *J* = 10.1, 3.5 Hz, 1H), 3.74 (s, 1H), 3.69 (d, *J* = 18.6 Hz, 1H), 3.35 (d, *J* = 18.6 Hz, 1H), 3.22 – 3.17 (m, 1H), 3.01 (dqd, *J* = 11.3, 7.3, 4.0 Hz, 1H), 2.63 (tdd, *J* = 13.6, 6.1, 1.8 Hz, 1H), 2.48 – 2.29 (m, 4H), 2.23 – 2.18(m, 2H), 2.14 – 2.09(m, 1H), 2.03 (td, *J* = 7.5, 2.7 Hz, 2H), 1.86 – 1.78 (m, 1H), 1.74 – 1.65 (m, 2H), 1.62 (t, *J* = 7.2 Hz, 2H), 1.57 (t, *J* = 7.2 Hz, 2H), 1.51 (s, 3H), 1.49 – 1.39 (m, 2H), 1.16 (ddd, *J* = 12.2, 8.1, 4.0 Hz, 1H), 0.93 (s, 3H), 0.82 (d, *J* = 7.3 Hz, 3H).

**^13^C NMR (125 MHz, CD_3_CN)** δ 211.35, 186.74, 167.97, 153.41, 130.02, 125.15, 101.97 (d, *J* = 174.4 Hz), 91.96, 84.01, 72.30 (d, *J* = 38.0 Hz), 70.24, 58.08, 49.12 (d, *J* = 22.7 Hz), 48.87, 44.73, 44.62, 44.60, 37.10, 36.30, 34.77 (d, *J* = 19.3 Hz), 33.53, 32.92, 32.87, 31.48, 28.24 (d, *J* = 4.1 Hz), 23.56 (d, *J* = 5.8 Hz), 17.32, 15.40, 13.62.

**HRMS (ESI)** m/z: [M+1]^+^ 512.2920, found 512.2917

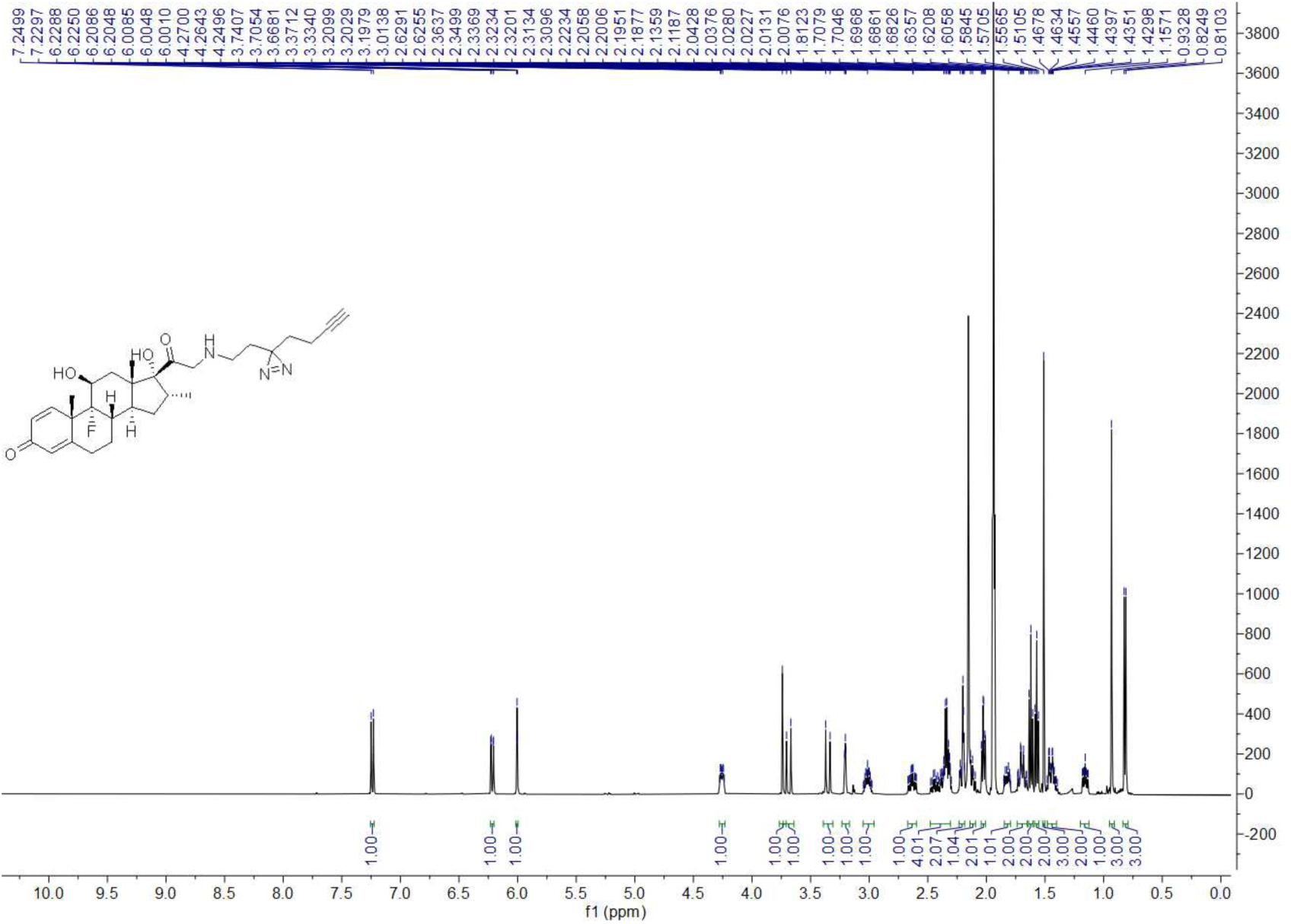

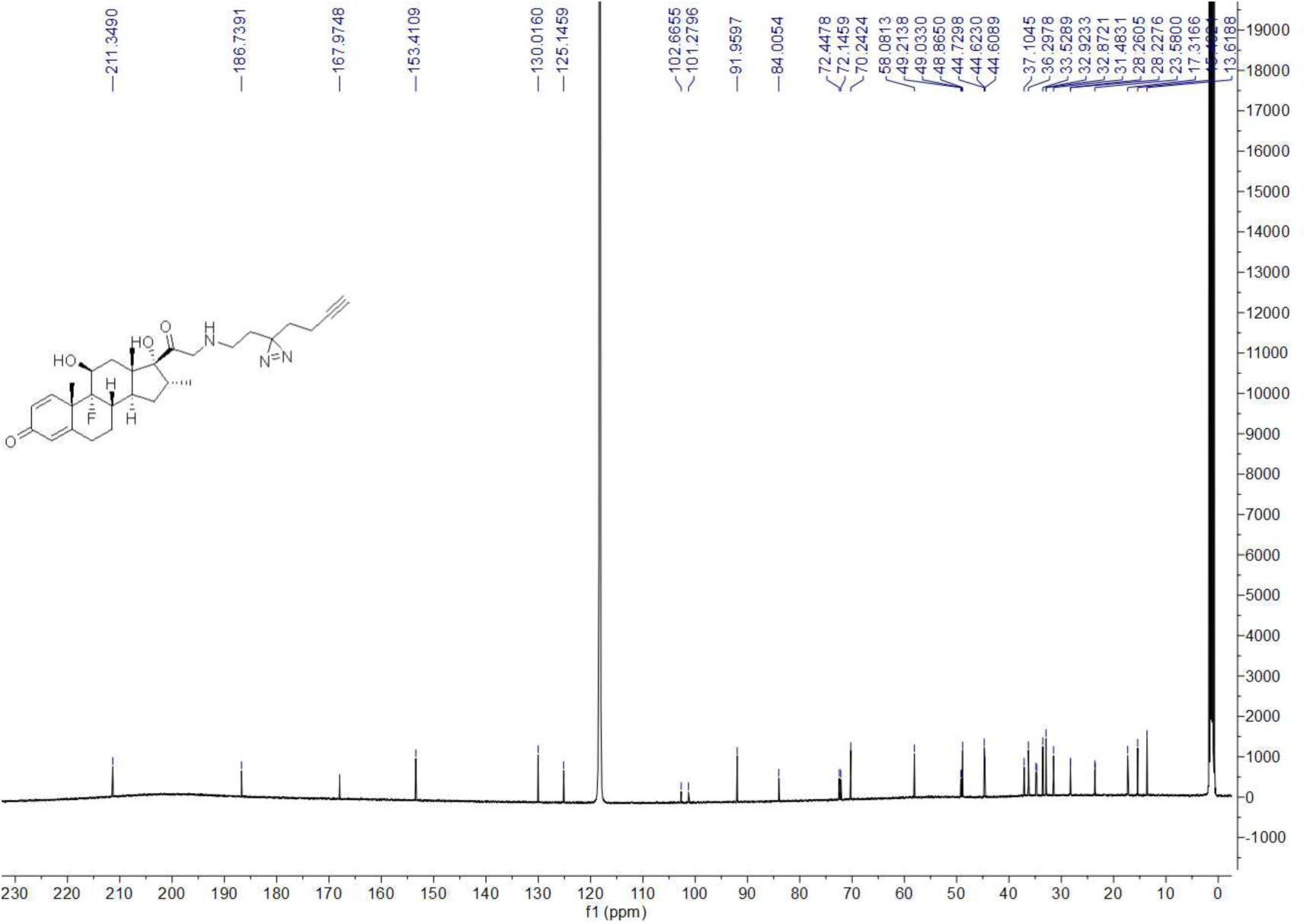

## In Situ Synthesis of TAMRA-NOR and TAMRA-DEX

TAMRA-NOR and TAMRA-DEX were synthesized in situ using copper-catalyzed azide-alkyne cycloaddition (CuAAC) chemistry. Briefly, the respective alkyne-functionalized probe was reacted with equimolar TAMRA-azide in the presence of 0.85 mM tris(benzyltriazolylmethyl)amine (TBTA), 1.2 mM copper(II) sulfate (CuSO₄), and 12 mM tris(2-carboxyethyl)phosphine (TCEP) for 90 min at room temperature under shaking conditions. The resulting reaction mixtures were used directly for microscopy experiments without further purification or workup.

## Data availability

PLACEHOLDER

## Author contributions

PLACEHOLDER

## Acknowledgements

PLACEHOLDER

---

**Supplementary Figure S1.**
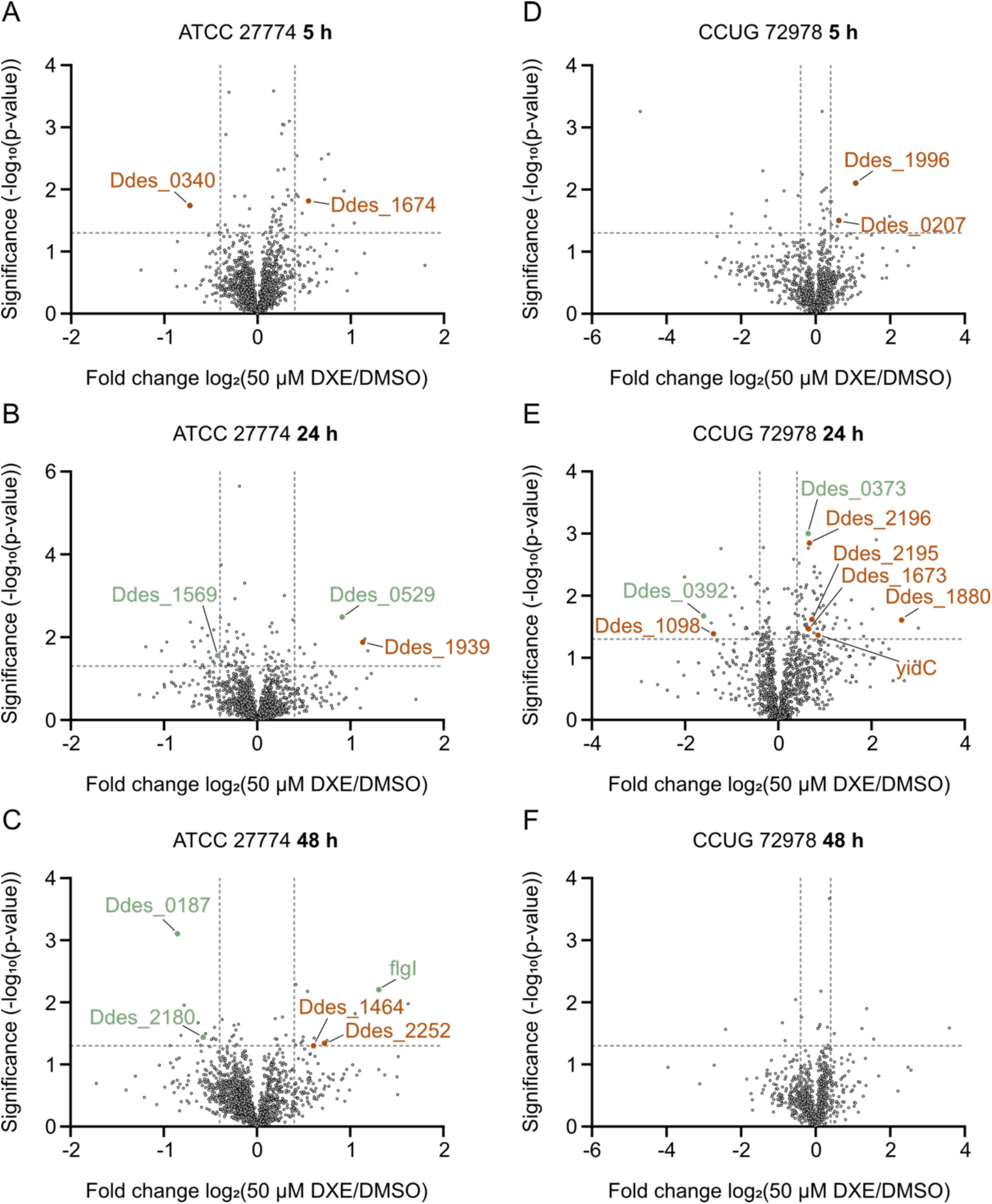
Differential abundance of flagella-associated proteins following DXE treatment in two *D. desulfuricans* strains. Volcano plots showing proteins associated with flagellar structure, assembly or regulation in *D. desulfuricans* ATCC 27774 (*A* – *C*) and CCUG 72978 (*D* – *F*) after 5 h, 24 h and 48 h incubation with **DXE**. Dashed vertical lines indicate fold-change thresholds of log₂FC = ±0.4, whereas the horizontal dashed line represents the significance threshold (−log₁₀ *p* = 1.3). Proteins directly involved in flagellar structure or assembly are highlighted in green, while proteins indirectly associated with flagellar function, motility or downstream regulatory pathways are shown in orange. Gene names are shown for significantly regulated proteins exceeding both thresholds that were classified as directly (green) or indirectly (orange) associated with flagellar processes.

**Supplementary Figure S2.**
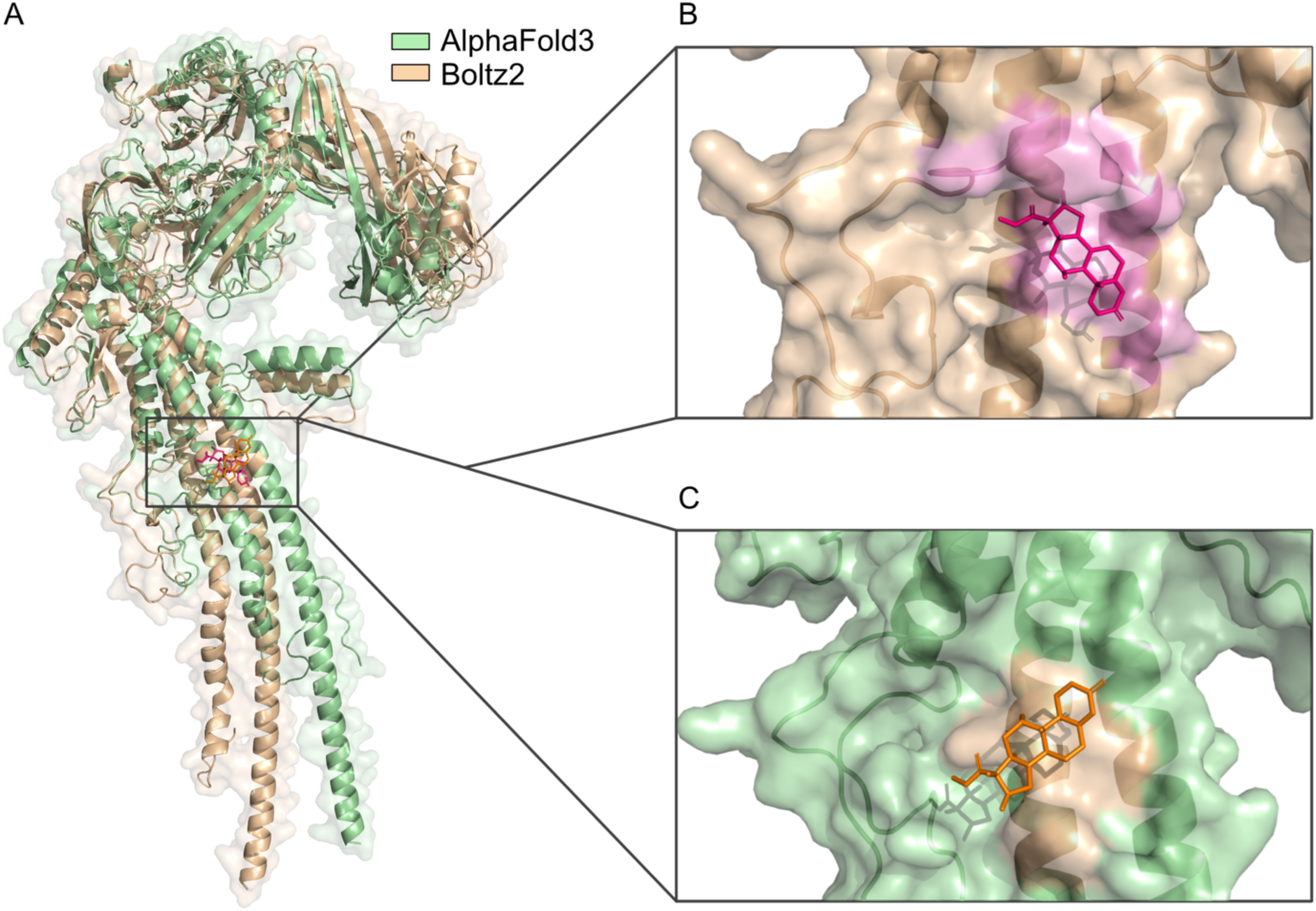
Comparison of AlphaFold3 and Boltz2 predictions for DXE binding to Ddes_0530. (*A*) Structural overlay of the AlphaFold3 and Boltz2 Ddes_0530-**DXE** complex predictions. The AlphaFold3 model is shown in green and the Boltz2 model in beige. Both methods localize **DXE** to the same D1-associated leg-region binding area. (*B*) Close-up of the Boltz2-predicted **DXE**-binding site. The Ddes_0530 surface is shown in beige and **DXE** is shown as magenta sticks, the local binding surface is highlighted in pink. (*C*) Close-up of the AlphaFold3-predicted **DXE**-binding site. The Ddes_0530 surface is shown in green and **DXE** is shown as orange sticks, the local binding surface is highlighted in beige.

**Supplementary Table S1.** Functional annotation of up-and downregulated proteins associated with flagellar structure, assembly, motility, and flagella-related regulatory pathways in *D. desulfuricans* ATCC 27774 and CCUG 72978 treated with DXE for 5, 24, or 48 h. Proteins were classified as either directly involved in flagellar structure and assembly (green) or indirectly associated with flagellar function through motility, signaling, membrane biogenesis, or energy metabolism (orange). Functional annotations are based on UniProt and InterPro database entries.

| Gene name | Protein name | UniProt ID | Regulation |
| --- | --- | --- | --- |
| **<u>ATCC 27774 5 h</u>** |  |  |  |
| ● Ddes_1674 | Putative diguanylate cyclase | B8J1E6 | up |
| ● Ddes_0340 | Na<sup>+</sup>/H<sup>+</sup> antiporter NhaC | B8J3L9 | down |
| **<u>ATCC 27774 24 h</u>** |  |  |  |
| ● Ddes_0529 | Flagellar protein FliS | B8J4I2 | up |
| ● Ddes_1939 | Putative diguanylate cyclase | B8J2R6 | up |
| ● Ddes_1569 | ATPase FliI/YscN | B8J143 | down |
| **<u>ATCC 27774 48 h</u>** |  |  |  |
| ● flgI | Flagellar P-ring protein | B8J2Y1 | up |
| ● Ddes_1464 | Sodium:proton antiporter | B8J0T9 | up |
| ● Ddes_2252 | ATP synthase protein I | B8J4L7 | up |
| ● Ddes_2180 | OmpA/MotB domain protein | B8J431 | down |
| ● Ddes_0187 | Two component, sigma54 specific, transcriptional regulator, Fis family | B8J2L7 | down |
| **<u>CUUG 72978 5 h</u>** |  |  |  |
| ● Ddes_0207 | MscS Mechanosensitive ion channel | B8J2N5 | up |
| ● Ddes_1996 | Outer membrane autotransporter barrel domain protein | B8J2X3 | up |
| **<u>CUUG 7297 24 h</u>** |  |  |  |
| ● Ddes_0373 | Flagellar protein FliL | B8J3Q1 | up |
| ● Ddes_1673 | NADH dehydrogenase (Quinone) | B8J1E5 | up |
| ● Ddes_1880 | NADH dehydrogenase (Quinone) | B8J2B7 | up |
| ● Ddes_2195 | Cytochrome d ubiquinol oxidase, subunit II | B8J446 | up |
| ● Ddes_2196 | Cytochrome bd ubiquinol oxidase, subunit I | B8J447 | up |
| ● yidC | Membrane protein insertase YidC | B8J2U4 | up |
| ● Ddes_0392 | Flagellar basal-body rod protein FlgG | B8J3S0 | down |
| ● Ddes_1098 | Lytic murein transglycosylase | B8IZS7 | down |
| **<u>CUUG 7297 48 h</u>** |  |  |  |
| - | - | - | - |

